# Post-translational modifications within fibrinaloid microclot complexes distinguish Pre-COVID-19 Postural Orthostatic Tachycardia Syndrome (POTS), Long COVID, and Long COVID-POTS and reveal disease-specific molecular pathways

**DOI:** 10.64898/2025.12.29.696828

**Authors:** Renata M Booyens, Mare Vlok, Cecile Bester, Rashmin Hira, M. Asad Khan, Douglas B. Kell, Satish R Raj, Etheresia Pretorius

## Abstract

**Background:** Pre-COVID-19 Postural orthostatic tachycardia syndrome (PC-POTS), Long COVID, and their overlap (LC-POTS) are chronic post-viral conditions marked by debilitating symptoms despite frequently normal routine laboratory results. We previously identified insoluble fibrinaloid microclot complexes (FMCs) in Long COVID. It is not known whether FMCs are also present in PC-POTS, or whether post-translational modifications (PTMs) within FMC-entrapped proteins contribute to disease mechanisms.

**Methods:** Platelet-poor plasma from healthy controls, PC-POTS patients (collected prior to the COVID-19 pandemic), Long COVID (without POTS) and LC-POTS patients underwent fluorescence imaging flow cytometry to quantify FMCs. Proteomic analyses were performed on insoluble protein fractions using a double trypsin digestion strategy and data-independent liquid chromatography-tandem mass spectrometry (LC-MS/MS). Differential protein abundance, PTMs, and amyloidogenicity were compared across groups.

**Results:** Measured with imaging flowcytometry in objects/mL, higher levels of FMCs were present in disease groups compared to controls, although not statistically significant. Statistically significant differences potentially lay within FMC sizes and composition. Furthermore, despite only a few dysregulated proteins, FMC proteomics revealed extensive and disease-specific peptides with PTM dysregulation across coagulation, immune, and metabolic pathways. Long COVID displayed FMCs with PTMs of coagulation proteins including prominent advanced glycation end-products (AGE)- and oxidation-based modifications of fibrinogen subunits, particularly fibrinogen subunit A (FIBA), resembling diabetic glycation profiles. FMCs in PC-POTS showed fewer fibrinogen PTMs but markedly increased modifications in metabolic proteins, including oxidised apoA1 and apoB, and immune patterns with complement-related proteins (C3, C4A/B, IC1), immunoglobulin G1 (IGG1) and alpha 2 macroglobulin (A2MG). LC-POTS shared coagulation pathology with Long COVID and immune pathology with PC-POTS. Many dysregulated peptides were determined by *in silco* methods to be highly amyloidogenic, consistent with FMCs as β-sheet-rich aggregates. Protein-level differences were minimal compared with PTM changes.

**Conclusions:** This study provides the first evidence that post-translational modifications (PTMs) within fibrinaloid microclots complexes (FMCs) uniquely distinguish pre-COVID-19 POTS, Long COVID, and Long COVID-POTS. Because PC-POTS samples were collected before SARS-CoV-2, their PTM patterns reflect intrinsic disease biology, allowing a clear separation from Long COVID-related changes. PTM profiling revealed pro-coagulant fibrinogen modifications in Long COVID and LC-POTS, metabolic-oxidative disruptions in Long COVID and PC-POTS, and immune dysregulation in PC-POTS and LC-POTS. None of these is detectable with routine assays, and all are independent of protein abundance. The consistent presence of amyloidogenic peptides suggests a contribution to microvascular dysfunction. These findings define disease-specific PTM landscapes and support new diagnostic and therapeutic avenues across autonomic and post-viral disorders.

## INTRODUCTION

Post Acute COVID-19 sequelae (PASC), or Long COVID, and postural orthostatic tachycardia syndrome (POTS) are chronic, debilitating, and primarily post-viral diseases ^1^. Long COVID is characterized by persistent symptoms after acute COVID-19 infections, affecting 10-26% of cases ^2,3^ across respiratory, systemic, musculoskeletal, neurological, cardiovascular, gastrointestinal, psychological, and dermatological categories ^2,4^. POTS involves autonomic dysfunction and an abnormal heart rate increase (>30 bpm) upon standing, predominantly in females (>90%), often causing light-headedness or fainting ^5–8^. Both Long COVID and POTS affect multiple physiological systems ^9–11^ but precise pathologies remain uncertain, and diagnoses are mostly symptom-based ^12^. Despite severe symptoms, routine blood analyses often show normal protein levels, leading clinicians to treat symptoms rather than underlying causes. In 2020, we identified dense amyloid-like deposits, Fibrinaloid Microclot Complexes (FMC), in acute COVID-19 and Type 2 Diabetes Mellitus (T2DM) patients, implicating coagulation dysfunction ^13^. In 2021, FMCs were observed in Long COVID blood, affecting the microcirculation ^14^. FMCs are aggregates of inflammatory proteins-mainly fibrinogen-in both normal and amyloid states, potentially explaining severe symptoms ^15^ whilst entrapping inflammatory molecules undetectable by conventional assays. FMCs also contain neutrophil extracellular traps (NETs) ^16^. Following the COVID-19 pandemic, a positive correlation between Long COVID and POTS incidences has been observed ^17–20^ suggesting shared pathophysiology and raising the possibility that pre-COVID-19 POTS (PC-POTS) and Long COVID POTS (LC-POTS) may exhibit FMCs ^21^. To our knowledge, FMCs in PC-POTS and LC-POTS have not yet been investigated, although it is known that many other diseases involving POTS and FMCs share comorbidities ^21^, and FMCs provide a mechanistically straightforward physiological explanation for POTS ^21^. Previously, we conducted proteomics on isolated FMCs in Long COVID to identify abundant proteins ^22^. Systematic reviews indicate that potential biomarkers reflect multiple mechanisms, and no concise profile supports a diagnostic criterion ^23^. Proteomics studies in POTS are limited, with mass-spectrometry analyses calling for further investigation of molecular mechanisms ^12^. Given that conventional protein analyses fail to fully explain disease pathophysiology, we investigated post-translational modifications (PTMs) as a potential mechanism. PTMs, enzymatic or non-enzymatic, can be reversible or irreversible; oxidative and metabolic stresses often induce irreversible non-enzymatic PTMs ^24^. PTMs on fibrinogen influence clot formation and degradation, contributing to pathological progression ^25^. Previous studies of Long COVID PTMs focused only on soluble plasma proteins, thereby missing PTMs of proteins entrapped in insoluble FMCs, potentially explaining severe symptoms despite normal lab values ^26^. We note, too, that the proteome of FMCs differs substantially from that of normal clots, being highly enriched in amyloid(ogenic) proteins ^27,28^; the same is true for ^29^ the macroclots observed in ischaemic stroke ^30^. We thus hypothesized that (1) FMCs are present in PC-POTS and LC-POTS, and (2) PTMs within FMCs should reveal disease-specific molecular changes that could reflect pathology even when conventional protein levels appear normal. To test this, our cohort included healthy controls, PC-POTS, Long COVID, and LC-POTS patients. FMC presence was measured by fluorescence imaging flow cytometry, followed by double trypsin digest isolation and liquid chromatography-tandem mass spectrometry (LC-MS/MS) proteomics. Analyses included protein profiling, biomarker discovery, and PTM evaluation.

## Methods and Materials

### Statistical Analyses

All statistical analyses were performed using Python (Version 3.14.2) with the SciPy library. Data visualization was created using Matplotlib and Seaborn packages.

### Flow Cytometry

FMC count (objects/mL) was analyzed across experimental groups. Descriptive statistics including sample size (N), mean, standard deviation, minimum, median, and maximum values were calculated for each group. Statistical analysis followed a structured four-step approach. First, a one-way analysis of variance (ANOVA) was performed to test for overall differences among groups. Second, residuals were extracted from the ANOVA model by calculating the difference between observed values and their respective group means. Third, the normality of residuals was assessed using the Shapiro-Wilk test (α = 0.05) to evaluate whether the ANOVA assumptions were met. Fourth, pairwise comparisons between all groups were conducted using the Mann-Whitney U test, a non-parametric alternative that does not assume normality. To control for multiple testing, Bonferroni correction was applied to the pairwise comparisons (adjusted α = 0.05/n, where n is the number of pairwise comparisons). Results were considered statistically significant after a Bonferroni correction at the adjusted α level, and nominally significant without correction at p < 0.05. Data visualization was performed using violin plot, box-and-whisker plots and individual data points overlaid.

### Mass-Spectrometry

For protein- and peptide level comparisons, fold changes were calculated between groups, and log2 fold change (log2FC) values were computed. Adjusted p-values were transformed to −log10(p-value) for volcano plot visualization. Significance thresholds were set at |log2FC| > 1 and adjusted p-value < 0.05. Features exceeding both thresholds were classified as significantly differentially abundant.

### Study participant details

#### Demographics table

Table 1 summarizes participant demographics.

**Table 1.**
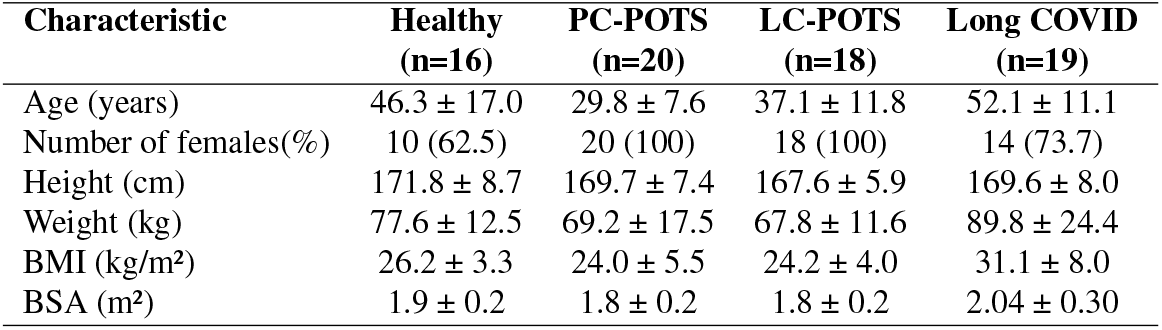
Participant Demographics: Data presented as mean ± SD or n (%). BMI: Body Mass Index; BSA: Body Surface Area; LC-POTS: Long COVID-Postural Orthostatic Tachycardia Syndrome; PC-POTS: Pre-COVID-19 Postural Orthostatic Tachycardia Syndrome.

| Characteristic | Healthy<br>(n=16) | PC-POTS<br>(n=20) | LC-POTS<br>(n=18) | Long COVID<br>(n=19) |
| --- | --- | --- | --- | --- |
| Age (years) | 46.3 ± 17.0 | 29.8 ± 7.6 | 37.1 ± 11.8 | 52.1 ± 11.1 |
| Number of females(%) | 10 (62.5) | 20 (100) | 18 (100) | 14 (73.7) |
| Height (cm) | 171.8 ± 8.7 | 169.7 ± 7.4 | 167.6 ± 5.9 | 169.6 ± 8.0 |
| Weight (kg) | 77.6 ± 12.5 | 69.2 ± 17.5 | 67.8 ± 11.6 | 89.8 ± 24.4 |
| BMI (kg/m<sup>2</sup>) | 26.2 ± 3.3 | 24.0 ± 5.5 | 24.2 ± 4.0 | 31.1 ± 8.0 |
| BSA (m<sup>2</sup>) | 1.9 ± 0.2 | 1.8 ± 0.2 | 1.8 ± 0.2 | 2.04 ± 0.30 |

#### Ethics and Informed Consent

Human samples used in this research were obtained from the University of Calgary where they had been collected under approvals from The Conjoint Health Research Ethics Board (CHREB), University of Calgary (REB21-1188 and REB15-2434). The transfer of samples from the University of Calgary to the University of Stellenbosch was conducted under a formal Material Transfer Agreement (MTA) (Contract number: S009247). The MTA governed the conditions of sample transfer, use, storage, and confidentiality, and confirmed that all materials were provided in accordance with the original ethics approval. All samples were received in de-identified form and were used solely for the purposes described in this study. No new samples were collected specifically for this study.

#### Study Participants and Diagnosis

Participants were recruited for studies focussed on Long COVID, POTS, and healthy controls. Participants whose samples were included in these analyses were adults aged 18-80 years of either sex. Long-COVID patients were required to have had a confirmed positive SARS-CoV-2 test and ongoing symptoms persisting for more than 12 weeks post-infection. The Long COVID cohort also presented with initial orthostatic hypotension ^31,32^. Healthy control participants had no history of confirmed or suspected SARS-CoV-2 infection, and no medical history consistent with POTS. POTS participants were adults aged 18-60 years, residing in Canada, with a physician-confirmed diagnosis according to the CCS Consensus Statement on Postural Orthostatic Tachycardia Syndrome ^33^, including sustained orthostatic tachycardia of ≥30 bpm within 10 minutes of standing without orthostatic hypotension, and chronic orthostatic symptoms improving with recumbence. Importantly, the Pre-COVID-19 POTS (PC-POTS) group was diagnosed before the COVID-19 pandemic and therefore had POTS unrelated to any COVID exposure. LC-POTS patients met criteria for Long-COVID and then later met the criteria for POTS. Exclusion criteria for all cohorts included-inability to provide informed consent, inability to safely withdraw from medications that could interfere with study assessments, and any other factors which, in the investigator’s opinion, would prevent completion of the study protocol, such as poor compliance or unpredictable scheduling. Participants needed the ability to attend the Calgary Autonomic Research Clinic for study procedures.

### Experimental Detail

#### Imaging Flow Cytometry

To establish the presence of FMCs in all groups, we conducted cell-free fluorescence imaging flow cytometry ^34^. PPP was stained with the fluorescent amyloid marker Thioflavin T (ThT). Images were processed using IDEAS^®^ 6.2 as developed for the Amnis^®^ FlowSight^®^ Imaging Flow Cytometer and total objects/mL were obtained.

#### Proteomics

A double trypsin digest was performed on platelet poor plasma (PPP) to isolate FMC. LC-MS/MS was used to identify and measure the proteins, peptides and PTMs present in the FMC using data independent acquisition (DIA). Detailed steps of the procedure have been reported in our previous studies ^22^. A group of proteins was selected for further analysis and included:

#### COAGULATION

- ANT3 (Antithrombin-III) - inhibits blood clotting
- FIBA, FIBB, FIBG (Fibrinogen alpha, beta, gamma chains) - forms fibrin clots
- ITA2B (Integrin alpha-IIb) - platelet aggregation
- MYH9 (Myosin-9) - platelet function and structure

#### IMMUNITY

- A2MG (Alpha-2-macroglobulin) - protease inhibitor, immune regulation
- CO3 (Complement C3) - central complement pathway protein
- CO4/B (Complement C4-A and C4-B) - complement activation
- IC1 (C1 inhibitor) - regulates complement and coagulation
- ITIH4 (Inter-alpha-trypsin inhibitor heavy chain H4) - acute phase response, protease inhibitor
- IGG1 (Immunoglobulin gamma-1) - antibody

#### METABOLIC

- APOA1 (Apolipoprotein A-I) - HDL cholesterol transport
- APOB (Apolipoprotein B) - LDL cholesterol transport
- ALBU (Albumin) - protein/lipid transport, osmotic pressure
- HBA, HBB (Hemoglobin alpha and beta) - oxygen transport
- ZA2G (Zinc-alpha-2-glycoprotein) - lipid mobilization

## RESULTS

### Imaging Flow Cytometry confirmed the presence of Fibrinaloid Microclot Complexes (FMC) in Postural Orthostatic Tachycardia Syndrome (POTS)

A one-way ANOVA was conducted to examine differences across the four groups. The overall effect was not statistically significant, (p = 0.14), indicating no significant differences in mean values among the groups. Following the ANOVA, residuals were extracted and tested for normality using the Shapiro-Wilk test. The test revealed that the residuals significantly deviated from normality, therefore non-parametric pairwise comparisons were conducted using Mann-Whitney U tests with Bonferroni correction for multiple comparisons (adjusted α = 0.0083).The pairwise comparisons revealed two nominally significant differences at the uncorrected alpha level: controls vs Long COVID (p = 0.011) and controls vs PC-POTS (p = .049). However, neither comparison remained statistically significant after applying the Bonferroni correction. All other pairwise comparisons were non-significant. These findings suggest that while the Long COVID and PC-POTS groups showed elevated values compared to controls, these differences did not reach the threshold for statistical significance when adjusting for multiple comparisons. Detailed descriptive statistics and pairwise comparison results are provided in Supplementary Tables S1.1,S1.2 and S1.3.

### Proteomic analysis of the insoluble FMC fraction revealed minimal protein differences

Proteins that were dysregulated in at least one disease group relative to controls were included in comparisons between disease groups. Only three proteins showed dysregulation at the protein concentration level (Figure 2). ZA2G was downregulated and MYH9 was upregulated in Long COVID compared to controls. Additionally, ZA2G was downregulated in Long COVID when compared to PC-POTS and LC-POTS (i.e. ZA2G is higher in PC-POTS and LC-POTS than in Long COVID). IC1 was downregulated in PC-POTS relative to controls. Detailed results are in Supplementary Table S2.

**Figure 1.**
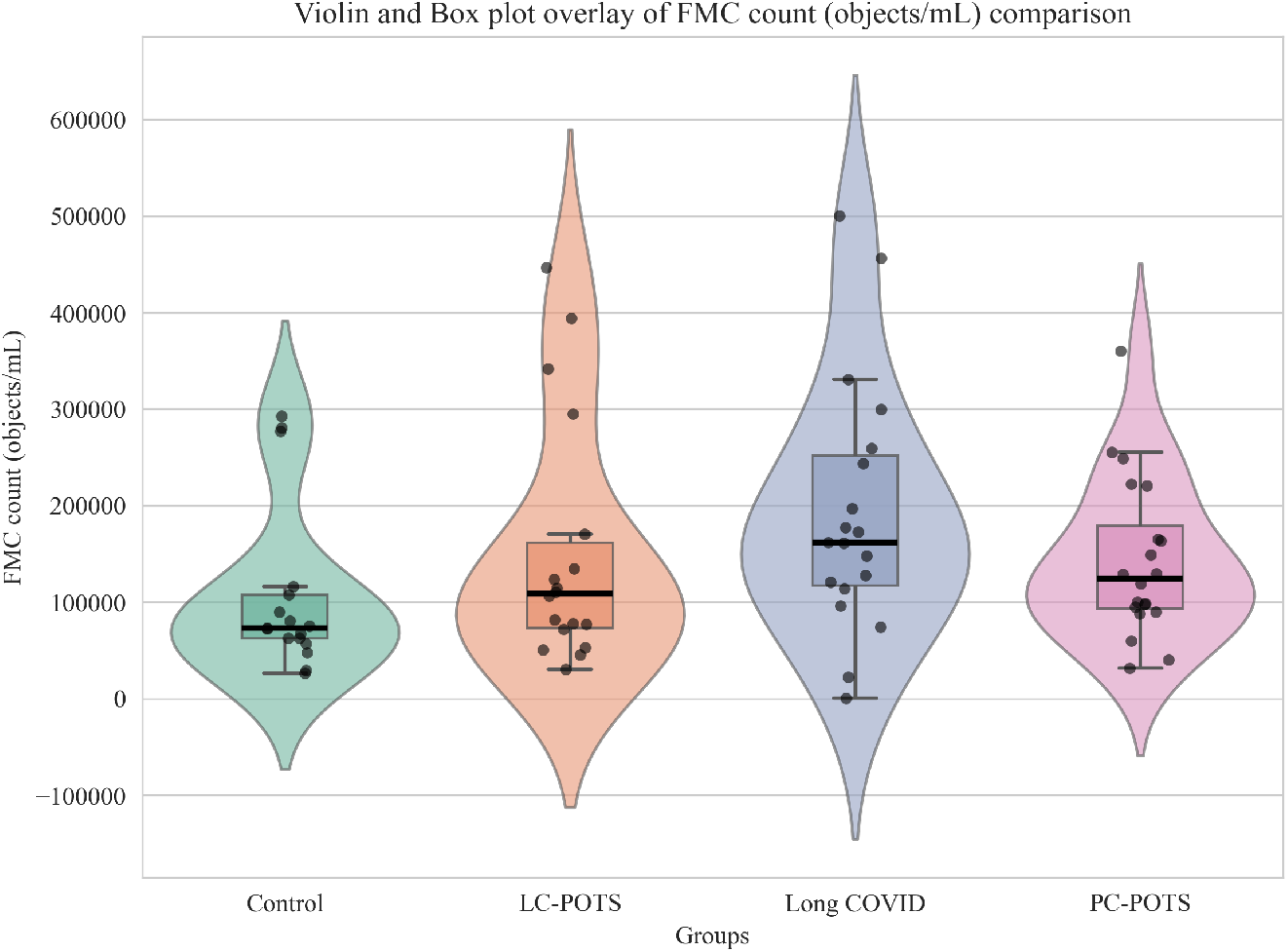
Box and Violin Plot Comparison of Fibrinaloid Microclot Complex (FMC) Counts. FMC counts (objects/mL) were measured across four groups: controls (*n* = 17, mean = 1.07 × 10^5^, SD = 8.73 × 10^4^), LC-POTS (*n* = 18, mean = 1.51 × 10^5^, SD = 1.28 × 10^5^), Long COVID (*n* = 19, mean = 1.93 × 10^5^, SD = 1.31 × 10^5^), and PC-POTS (*n* = 20, mean = 1.43 × 10^5^, SD = 8.25 × 10^4^). A one-way ANOVA revealed no statistically significant overall difference among groups (*p* = 0.14). Due to significant deviation from normality in the ANOVA residuals (Shapiro-Wilk test, *p <* 0.001), non-parametric pairwise comparisons were conducted using Mann-Whitney U tests with Bonferroni correction (adjusted *α* = 0.0083). Two comparisons showed nominal significance at the uncorrected *α* = 0.05 level: controls vs Long COVID (*p* = 0.011) and controls vs PC-POTS (*p* = 0.049); however, neither remained significant after Bonferroni correction. Box plots display median (center line), interquartile range (box), and range (whiskers), while violin plots show the full distribution of data points for each group. POTS: Postural Orthostatic Tachycardia Syndrome; PC: Pre-COVID; LC: Long COVID.

**Figure 2.**
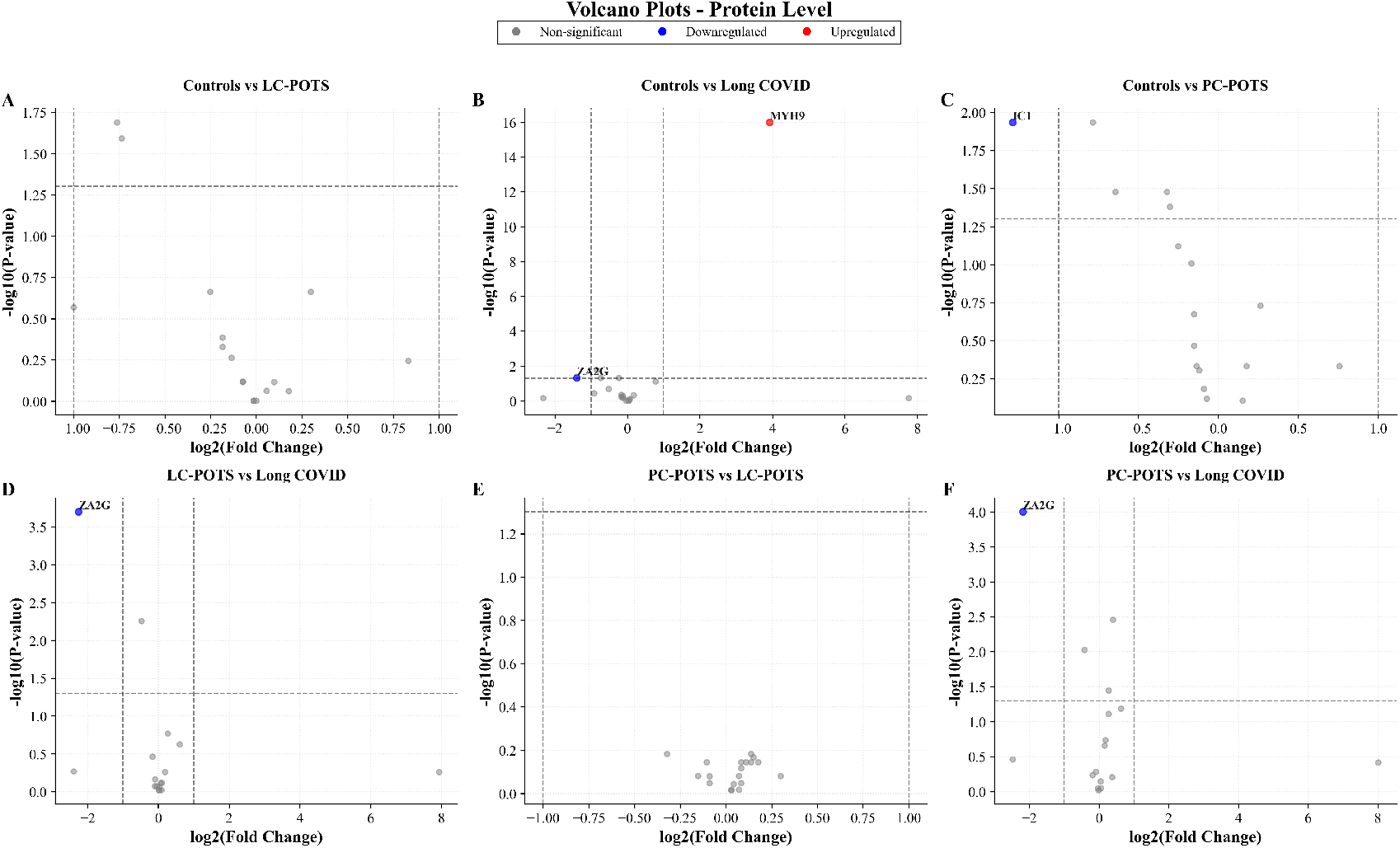
Volcano Plots Illustrating Differential Protein Regulation Across Pairwise Group Comparisons. For protein-level comparisons, fold changes were calculated between groups, and log_2_ fold change (log_2_FC) values were computed. Adjusted *p*-values were transformed to − log_10_(*p*-value) for visualization. The *x*-axis represents log_2_ fold change, and the *y*-axis represents − log_10_(*p*-value). Significance thresholds were set at | log_2_ FC| *>* 1 and adjusted *p*-value *<* 0.05. Features exceeding both thresholds were classified as significantly differentially abundant. Horizontal dotted lines indicate the −log_10_(*p*-value) cutoff, and vertical dotted lines indicate the log_2_ fold change cutoff (±1). **A)** Controls vs. LC-POTS: No proteins were significantly different. **B)** Controls vs. Long COVID: Two proteins were significantly different (MYH9 upregulated; ZA2G downregulated). **C)** Controls vs. PC-POTS: Only IC1 was significantly downregulated. **D)** LC-POTS vs. Long COVID: ZA2G was significantly downregulated. **E)** PC-POTS vs. LC-POTS: No proteins were significantly different. **F)** PC-POTS vs. Long COVID: ZA2G was significantly downregulated.POTS: Postural Orthostatic Tachycardia Syndrome, PC: Pre-COVID; LC: Long COVID

### There is Extensive PTM dysregulation within the insoluble FMC fraction

The proteomics analysis which included post translational modifications revealed 29 unique base peptides which showed significant differences between controls and at least 1 disease group - 58.62% of the base peptides were amyloidogenic according to Amylogram (results in Supplementary Table S3.1, and S3.2), i.e. had AmyloGram scores greater than 0.5. Furthermore, these 29 base peptides originating from 16 proteins, included 64 uniquely modified peptides and revealed multiple significant differences between groups (Figure 3 and Table 2). Detailed results are available in Supplementary Tables S4.1, S4.2 and S4.3.

**Table 2.** Results from pair wise peptide and PTM comparison. For all comparisons (X vs Y), regulation is defined relative to the second group (Y). Up-regulation and down-regulation therefore reflect increased or decreased levels in Y compared with X, respectively.* Total base peptides are those peptides that were modified and whose modified, or unmodified, forms showed differences between groups. **Total proteins are the proteins represented by these base peptides. ***Total up- and downregulated counts refer to both the modified and unmodified forms of peptides that differed between groups.

| Category | Comparison | Total Proteins** | Total Base Peptide Seq.* | Total Dys-regulated Modified | Total Dys-regulated Unmodified | Total Up-regulated*** | Total Down-regulated*** |
| --- | --- | --- | --- | --- | --- | --- | --- |
| Coagulation | Controls vs Long COVID | 2 | 2 | 2 | 0 | 2 | 0 |
|  | Controls vs LC-POTS | 1 | 1 | 1 | 0 | 1 | 0 |
|  | Controls vs PC-POTS | 3 | 4 | 16 | 0 | 2 | 14 |
|  | LC-POTS vs Long COVID | 3 | 3 | 3 | 0 | 1 | 2 |
|  | PC-POTS vs Long COVID | 3 | 4 | 21 | 0 | 19 | 2 |
|  | PC-POTS vs LC-POTS | 0 | 0 | 0 | 0 | 0 | 0 |
| Immunity | Controls vs Long COVID | 1 | 2 | 2 | 0 | 1 | 1 |
|  | Controls vs LC-POTS | 3 | 5 | 9 | 0 | 7 | 2 |
|  | Controls vs PC-POTS | 6 | 12 | 20 | 0 | 13 | 7 |
|  | LC-POTS vs Long COVID | 3 | 6 | 14 | 0 | 2 | 12 |
|  | PC-POTS vs Long COVID | 3 | 6 | 14 | 0 | 1 | 13 |
|  | PC-POTS vs LC-POTS | 1 | 1 | 1 | 0 | 0 | 1 |
| Metabolic | Controls vs Long COVID | 3 | 3 | 3 | 0 | 0 | 3 |
|  | Controls vs LC-POTS | 2 | 2 | 5 | 0 | 4 | 1 |
|  | Controls vs PC-POTS | 3 | 6 | 8 | 1 | 5 | 4 |
|  | LC-POTS vs Long COVID | 2 | 2 | 5 | 0 | 0 | 5 |
|  | PC-POTS vs Long COVID | 2 | 4 | 6 | 1 | 2 | 5 |
|  | PC-POTS vs LC-POTS | 0 | 0 | 0 | 0 | 0 | 0 |

**Figure 3.**
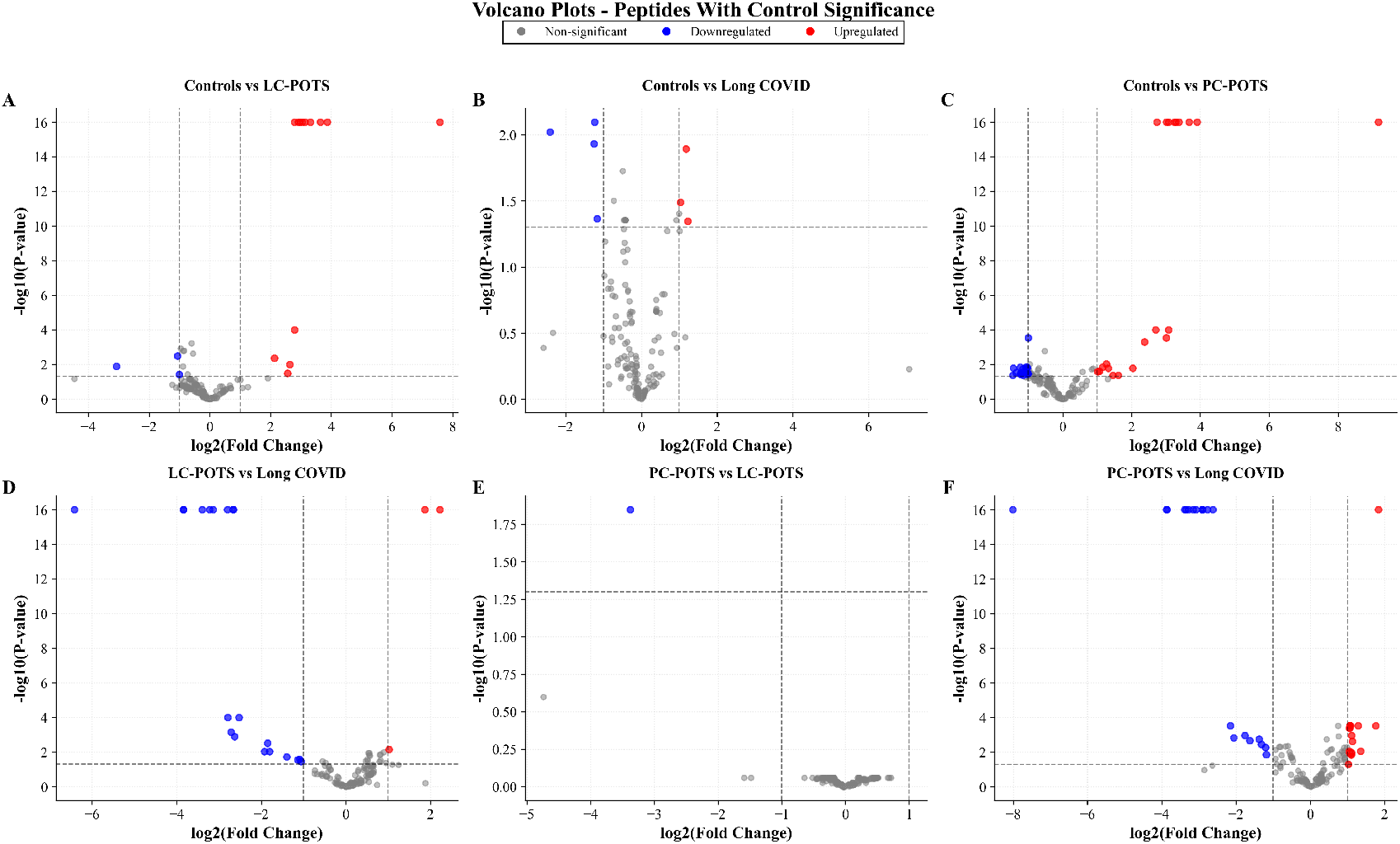
Volcano Plots Illustrating Differential Peptides with Post-Translational Modifications (PTM) Across Pairwise Group Comparisons. For peptide-level comparisons, pre-calculated log_2_ fold change values from mass spectrometry data were used. Adjusted *p*-values were transformed to − log_10_(*p*-value) for visualization. The *x*-axis represents log_2_ fold change, and the *y*-axis represents − log_10_(*p*-value). Significance thresholds were set at | log_2_ FC| *>* 1 and adjusted *p*-value *<* 0.05. Features exceeding both thresholds were classified as significantly differentially abundant. Horizontal dotted lines indicate the −log_10_(*p*-value) cutoff, and vertical dotted lines indicate the log_2_ fold change cutoff (±1). Analysis of 29 unique base peptides and 64 uniquely modified peptides across 16 proteins revealed several significant differences between groups. **A)** Controls vs. LC-POTS: 15 uniquely modified peptides (8 base peptides from 6 proteins) were dysregulated (12 upregulated, 3 downregulated). **B)** Controls vs. Long COVID: 7 uniquely modified peptides (7 base peptides from 6 proteins) were dysregulated (3 upregulated, 4 downregulated). **C)** Controls vs. PC-POTS: 44 uniquely modified peptides and one unmodified peptide (22 base peptides from 12 proteins) were dysregulated (20 upregulated, 25 downregulated). **D)** LC-POTS vs. Long COVID: 22 uniquely modified peptides (11 base peptides from 8 proteins) were significantly different (3 upregulated, 19 downregulated). **E)** PC-POTS vs. LC-POTS: 1 downregulated modified peptide. **F)** PC-POTS vs. Long COVID: 41 uniquely modified peptides and one unmodified peptide (14 base peptides from 8 proteins) were dysregulated (22 upregulated, 20 downregulated). POTS: Postural Orthostatic Tachycardia Syndrome, PC: Pre-COVID; LC: Long COVID

## DISCUSSION

In this study, we investigated whether FMCs are present in PC-POTS and if their associated peptides with post-translational modifications (PTM) distinguish PC-POTS, Long COVID and LC-POTS at the molecular level. The PC-POTS samples were collected before the COVID-19 pandemic, enabling us to define the baseline molecular signatures of classic POTS entirely independent of SARS-CoV-2 exposure. The LC-POTS group were all diagnosed with POTS after an acute COVID-19 infection. The pre-pandemic POTS cohort therefore provides a true reference for classical POTS biology, allowing us to distinguish molecular features that originate from established POTS mechanisms from those newly introduced or amplified by Long COVID in individuals who later developed POTS symptoms. Imaging flow cytometry confirmed the presence of FMCs across all disease groups, indicating that FMC formation is not limited to Long COVID. Whilst total FMC counts did not differ significantly between cohorts (after Bonferroni correction), the trend toward higher FMC counts in disease groups underscores the limitations of relying solely on FMC quantity and highlights the importance of examining detailed FMC composition, through proteomics. Proteomic analysis of the insoluble FMC fraction revealed that widespread dysregulation of PTMs far exceeded changes observed at the level of protein abundance. This suggests that PTMs, rather than protein concentration, more accurately reflect the underlying disease biology.

### Coagulation-related PTMs and FMC structural biology

Several fibrinogen peptides displayed disease-specific PTM patterns, underscoring the central role of coagulation proteins in FMC composition. In Long COVID, increased advanced glycation end-products (AGE) modifications on FIBA peptides align with known glycation patterns observed in diabetes ^35^. Furthermore, AGEs results in the formation of covalent crosslinks, which may help explain the robust, fibrinolysis-resistant nature of FMCs in this group ^36^. Conversely, PC-POTS showed reduced modification of selected FIBG peptides, which may reflect structural shielding of these sites once fibrinogen is incorporated into compact FMCs ^37^. Oxidation, deamination and pyroglutamate (PYD) formation of fibrinogen subunits collectively suggest altered clot architecture across diseases, with implications for fibrinolysis, clot permeability, and microvascular obstruction, which was also seen in the LC-POTS group. The presence of amyloidogenic peptides in multiple fibrinogen regions supports the concept that FMCs arise from misfolded, β-sheet-rich aggregates rather than classical thrombin-driven fibrin ^38^. At the protein level, MYH9 is upregulated in Long COVID alongside peptides of ITA2B, which exhibit post-translational modification (PTM) changes indicative of dysregulated anticoagulant function and could allude to altered platelet-fibrinogen interactions ^39,40^. The latter could also apply to the PC-POTS group as dysregulation of ANT3 peptides was observed within that group. Collectively, these findings indicate that the coagulation system is particularly disrupted in Long COVID and LC-POTS (Figure 4), with FMC-associated PTMs potentially amplifying microvascular dysfunction.

**Figure 4.**
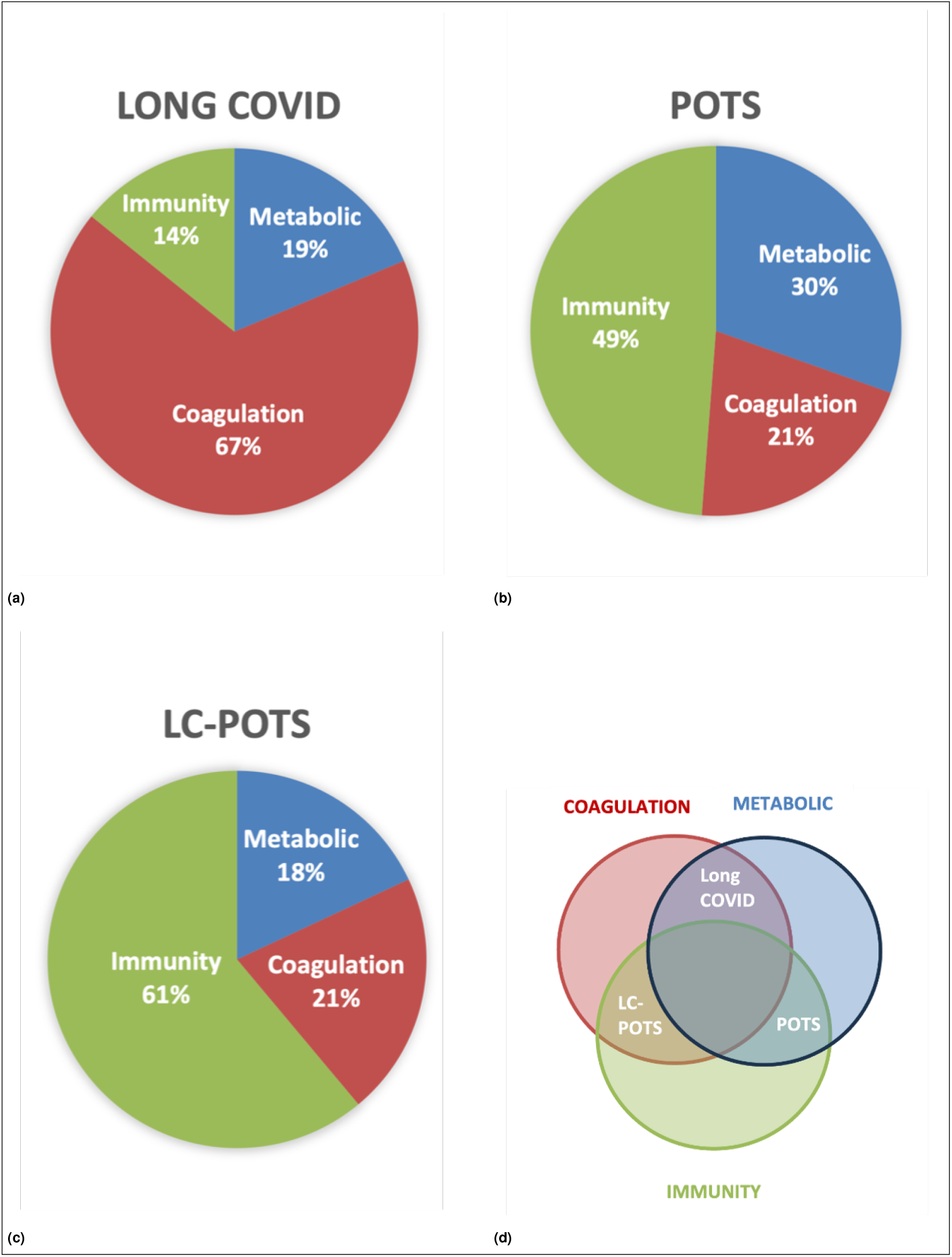
Functional category contribution of dysregulated peptides across disease groups. **a-c)** Pie charts illustrate the relative contribution of dysregulated peptides and proteins to three functional categories (metabolic, coagulation, and immunity) within each disease group. For each group, log_2_ fold-change values were calculated relative to controls and from pairwise comparisons between disease groups, and peptides were included according to the direction of regulation consistent with the expected biological effect of disease on the respective pathways. Log_2_ fold-change values of dysregulated peptides and proteins were summed within each functional category to generate a relative measure of category contribution, which was then expressed as a percentage of the total summed signal per disease group. These values are intended to provide a visual summary of dominant biological themes and were not used for statistical inference. **d)** A schematic Venn diagram summarizes the dominant functional categories across disease groups based on the two highest-contributing categories per group, highlighting conceptual overlap in biological processes rather than shared peptides or proteins.

### Immune and complement system dysregulation

PTMs in proteins involved in complement activation and innate immunity further differentiate the disease groups. C3, C4A/B and IGG1 peptides displayed distinct PTM profiles in PC-POTS compared to controls, suggesting altered complement turnover and immune activation ^41^. Downregulation of C1 inhibitor (IC1) in PC-POTS parallels patterns seen in complement/contact activation disorders such as hereditary angioedema, which share clinical overlap with POTS, including hypotension, autonomic instability and inflammatory symptoms. Additionally, the complement system plays a central role in mast cell activation and allergic inflammation ^42^, and several clinical studies have reported substantial overlap between mast cell activation syndrome (MCAS) and POTS ^43^. A2MG showed multiple oxidation- and deamination-based PTMs, particularly in PC-POTS and LC-POTS. Given A2MG’s role in trapping proteases and binding cytokines, these modifications could alter inflammatory signalling ^44,45^. The pattern observed for ITIH4 (reduced oxidised forms in PC-POTS despite its identity as an acute-phase reactant) may indicate an oxidative environment that selectively preserves functional ITIH4, potentially contributing to sustained inflammatory signalling ^46^. Collectively, these findings suggest distinct immune-modulatory landscapes across diseases, especially in PC-POTS and LC-POTS groups.

### Metabolic pathway dysregulation

PTMs in lipid-handling proteins were prominent in PC-POTS and LC-POTS. Oxidised and deaminated variants of apoA1 and apoB were consistently upregulated in PC-POTS relative to both controls and Long COVID. This pattern resembles PTM signatures observed in T2DM, where oxidised apolipoproteins contribute to vascular dysfunction and altered lipid metabolism ^47,48^. ZA2G downregulation in PC-POTS and Long COVID aligns with insulin resistance pathways and supports a metabolic contribution to symptoms ^49,50^. These findings indicate that PC-POTS is characterized by a pronounced metabolic-oxidative PTM profile. Moreover, because the metabolic dysregulation observed in Long COVID - specifically AGE- and oxidation-mediated modifications of fibrinogen - mirrors that seen in T2DM, the Long COVID group likewise exhibits dysregulation of metabolic pathways (Figure 4).

### Cross-disease integration and mechanistic interpretation

Synthesising these data, Long COVID appears to be predominantly characterised by PTM dysregulation of coagulation proteins, particularly fibrinogen, with downstream effects on both coagulation and metabolic pathways. In contrast, both PC-POTS and LC-POTS exhibit more pronounced PTM alterations within immune pathways; PC-POTS shows secondary involvement of metabolic modifications, whereas LC-POTS demonstrates secondary dysregulation of coagulation pathways. The increase in AGE-modified FIBA peptides in Long COVID mirrors glycation patterns in diabetes; a convergence with notable clinical relevance, given that metformin reduces the risk of developing Long COVID ^51^. The downregulation of FIBG modifications in PC-POTS suggests structural differences in FMC architecture that may shield specific fibrinogen sites from modification. The high amyloidogenicity of many dysregulated peptides further reinforces the nature of FMCs as aggregated, amyloid-prone complexes. Importantly, the differential modification of individual peptides within the same protein indicates site-specific PTM non-enzymatic modifications rather than uniform protein-level changes, offering a molecular explanation for why routine soluble plasma assays often fail to capture disease pathology.

### Clinical relevance and implications for biomarker development

Samples from Long COVID patients exhibited widespread PTM alterations in coagulation-related proteins, particularly AGE- and oxidation-based modifications of fibrinogen subunits, reflecting a strong pro-coagulant signature, whereas PC-POTS presented a predominantly metabolic-oxidative PTM pattern affecting apolipoproteins and pointing to metabolic dysregulation. Secondarily PC-POTS affects the immune system through IGG1 and complement systems proteins. Interestingly, both conditions share features commonly associated with diabetes, such as protein glycation and oxidative stress. LC-POTS shares features with Long COVID in the coagulation pathway, and with PC-POTS in immunity (Figure 4). Together, these findings highlight PTMs within FMCs as a mechanistically rich and clinically relevant domain of investigation. By analysing the insoluble fraction, we have uncovered biochemical perturbations invisible to standard blood tests, offering a potential explanation for persistent and debilitating symptoms despite “normal” laboratory values. PTM signatures, especially those involving fibrinogen, complement proteins, and apolipoproteins, may hold utility as diagnostic or stratification biomarkers, and the distinct FMC-associated PTM patterns observed across Long COVID, PC-POTS and LC-POTS could inform disease-specific therapeutic strategies, including interventions targeting glycation pathways, oxidative stress, or complement activation

### Limitations and future directions

This study focused exclusively on the insoluble FMC fraction; parallel analysis of soluble proteins would clarify how PTMs partition between fractions. Larger cohorts are needed to validate disease-specific PTM signatures and to correlate molecular findings with symptom severity. Functional assays are required to determine how individual PTMs affect protein activity, fibrinogen structure, or complement regulation. Including a T2DM cohort in future studies will be essential, given the strong overlap between Long COVID and diabetic PTM patterns. Future work should also examine size-resolved FMC characteristics using advanced imaging flow cytometry to capture structural differences that may be clinically meaningful.

## Conclusion

This study provides the first direct evidence that PMTs within insoluble FMCs distinctly characterize PC-POTS, Long COVID, and LC-POTS. Since the PC-POTS samples predate the COVID-19 pandemic, the PTM patterns represent intrinsic POTS features rather than SARS-CoV-2 consequences, enabling the contributions of classical POTS biology and Long COVID dysregulation to be disentangled. PTM profiling reveals distinct yet overlapping biochemical signatures invisible to standard blood tests: Long COVID exhibits pro-coagulant fibrinogen modifications with metabolic dysregulation. PC-POTS and LC-POTS shows predominantly immune-oxidative disruptions, with POTS also showing metabolic dysregulation and LC-POTS include oxidation-modifications effecting coagulation. These PTM landscapes highlight potential avenues for biomarker-based diagnosis, patient stratification, and therapeutic strategies targeting glycation, oxidative stress, and complement pathways. The consistent presence of amyloidogenic peptides across groups underscores FMCs as β-sheet-rich, fibrinolysis-resistant aggregates contributing to microvascular dysfunction. Minimal protein-abundance differences alongside substantial PTM dysregulation provide a mechanistic explanation for severe symptoms despite normal routine lab findings. Overall, these findings offer a unifying framework and the first provable hypothesis for shared and divergent mechanisms in post-viral and dysautonomic conditions, establishing a foundation for biomarker-driven diagnostics and personalized therapeutics across the spectrum of autonomic and post-viral disorders.

## ACKNOWLEDGMENTS

This work was funded by StandingUpToPOTS 2024/2025 Pretorius, Kell and Lip grant.

## AUTHOR CONTRIBUTIONS

**Methodology:** RB, EP, MV, SRR, RH **Sample acquisition:** RH, SRR **Investigation:** RB, EP, MV **Visualisation and analyses:** RB, CB, MV **Funding acquisition:** EP, DK **Project administration:** RB, MV, SRR, RH **Writing original draft:** RB, EP **Writing, review and editing:** RB, EP, CB, MV, SR, RH, DK, AK

## RESOURCE AVAILABILITY

### Lead contact

Requests for further information and resources should be directed to and will be fulfilled by the lead contact, Etheresia Pretorius and Renata M. Booyens

### Materials and Data Availability

This study did not generate new materials. Any additional information required to reanalyse the data reported in this paper is available from the lead contact upon request.

## Supplementary Materials

**Table S1.1.** Descriptive Statistics: Fibrinaloid Microclot Complexes (objects/mL)

| Group | N | Mean | Std Dev | Min | Median | Max |
| --- | --- | --- | --- | --- | --- | --- |
| Control | 17 | 107,047 | 87,347 | 26,234 | 73,374 | 292,575 |
| LC-POTS | 18 | 151,462 | 127,683 | 30,736 | 108,659 | 446,460 |
| Long COVID | 19 | 192,916 | 131,474 | 539 | 161,950 | 500,164 |
| PC-POTS | 20 | 143,256 | 82,461 | 31,737 | 124,156 | 360,446 |

**Table S1.2.**
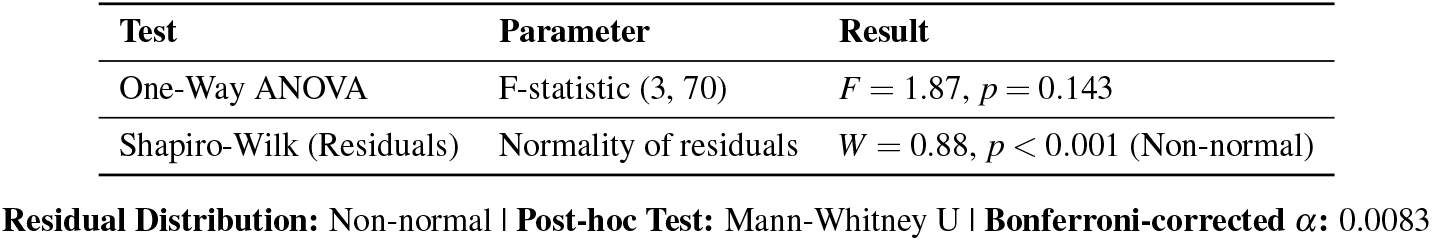
Statistical Tests Summary.

**Table S1.3.** Pairwise Comparisons with Bonferroni Correction.

| Comparison | Test | p-value | Significance |
| --- | --- | --- | --- |
| Control vs LC-POTS | Mann-Whitney U | 0.1607 | ns |
| Control vs Long COVID | Mann-Whitney U | 0.0112 | * |
| Control vs PC-POTS | Mann-Whitney U | 0.0493 | * |
| LC-POTS vs Long COVID | Mann-Whitney U | 0.1325 | ns |
| LC-POTS vs PC-POTS | Mann-Whitney U | 0.5686 | ns |
| Long COVID vs PC-POTS | Mann-Whitney U | 0.2112 | ns |

**Significance Legend:** \*\*\* = Significant after Bonferroni correction ( $p < 0.0083$ ); \* = Significant without correction ( $p < 0.05$ ); ns = Not significant

### Supplementary Table 2: Protein Dysregulation Proteomics Results

**Table S2.**
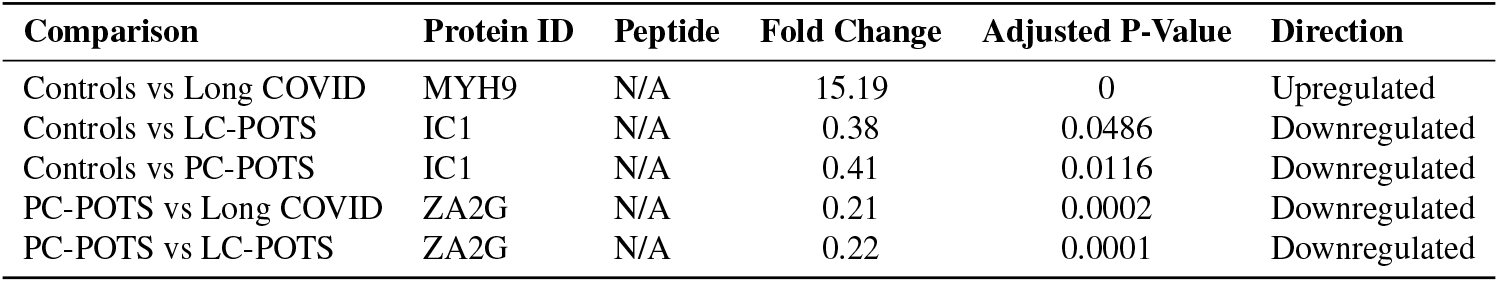
Proteomics Protein Level Analyses.

### Supplementary Table 3: Amylogram results

**Table S3.1.** Amyloidogenic Prediction Results for Peptide Sequences.

| Sequence | Score | Amyloidogenic |
| --- | --- | --- |
| AIQLTYNPDESSKPNMIDAATLK | 0.667 | Yes |
| ASVQLLVQDSLNPVK | 0.890 | Yes |
| DDNPNLPRLVRPEVDVMCTAFHDNEETFLK | 0.535 | Yes |
| DLGEENFK | 0.078 | No |
| EFNAETFTFHADICTLSEKER | 0.410 | No |
| FNWYVDGVEVHNAK | 0.705 | Yes |
| FQPTLLTLPR | 0.367 | No |
| IAQWQSFQLEGGLK | 0.468 | No |
| IQKLESDVSAQMEYCR | 0.110 | No |
| LLDNWDSVTSTFSK | 0.568 | Yes |
| LLVVYPWTQR | 0.726 | Yes |
| LVNGQSHISLSK | 0.666 | Yes |
| MFLSFPTTK | 0.455 | No |
| NMEQFQVSVSVAPNAK | 0.511 | Yes |
| NQVSLTCLVK | 0.507 | Yes |
| NTLYLNMNSLRPEDTAVYYCAR | 0.755 | Yes |
| PKDTLMISRTPEVTCVVVDVSHEDPEVK | 0.667 | Yes |
| PLLEKSHCIAEVENDEMPADLPSLAADFVESK | 0.516 | Yes |
| QFSFPLSSEPFQGSYK | 0.324 | No |
| RLEVDIDIK | 0.408 | No |
| RPCFSALEVDETYVPK | 0.666 | Yes |
| SEETKENEGFTVTAEGK | 0.327 | No |
| THTCPPCPAPELLGGPSVFLFPPKPK | 0.524 | Yes |
| VAEGTQVLELPFKGDDITMVLILPKPEK | 0.699 | Yes |
| VELEDWNGR | 0.273 | No |
| VVSVLTVLHQDWLNGK | 0.669 | Yes |
| WLNEQRYYYGGYGSTQATFMVFQALAQYQK | 0.457 | No |
| YENYELTLK | 0.511 | Yes |
| YYGGGYGSTQATFMVFQALAQYQK | 0.457 | No |

**Table S3.2.** Summary Statistics.

| Metric | Value |
| --- | --- |
| Total Peptides | 29 |
| Amyloidogenic Peptides | 17 |
| Non-Amyloidogenic Peptides | 12 |
| Percentage Amyloidogenic | 58.62% |

### Supplementary Table 4: Peptide and PTM Dysregulation Proteomics Results

**Table S4.1.**
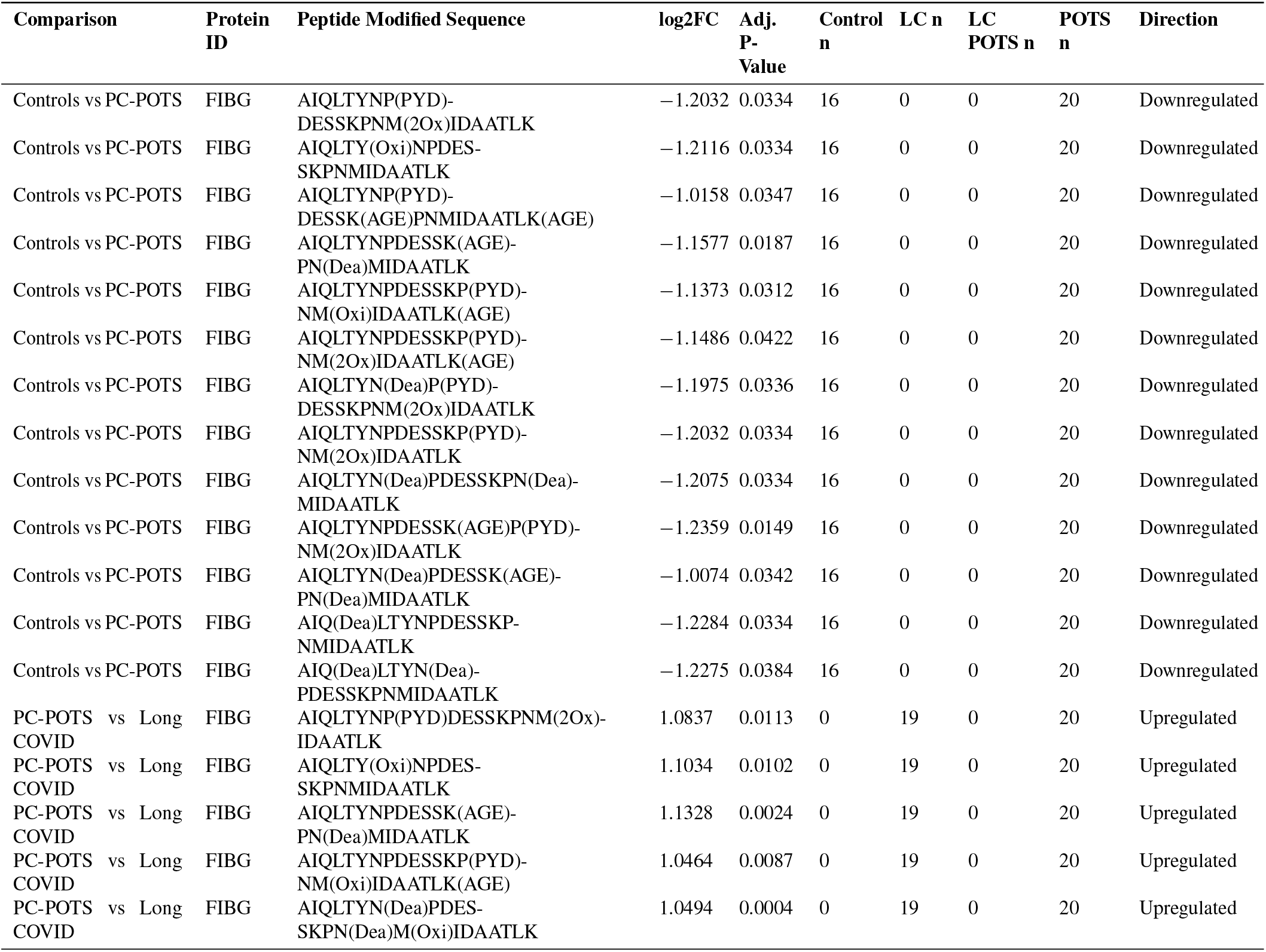
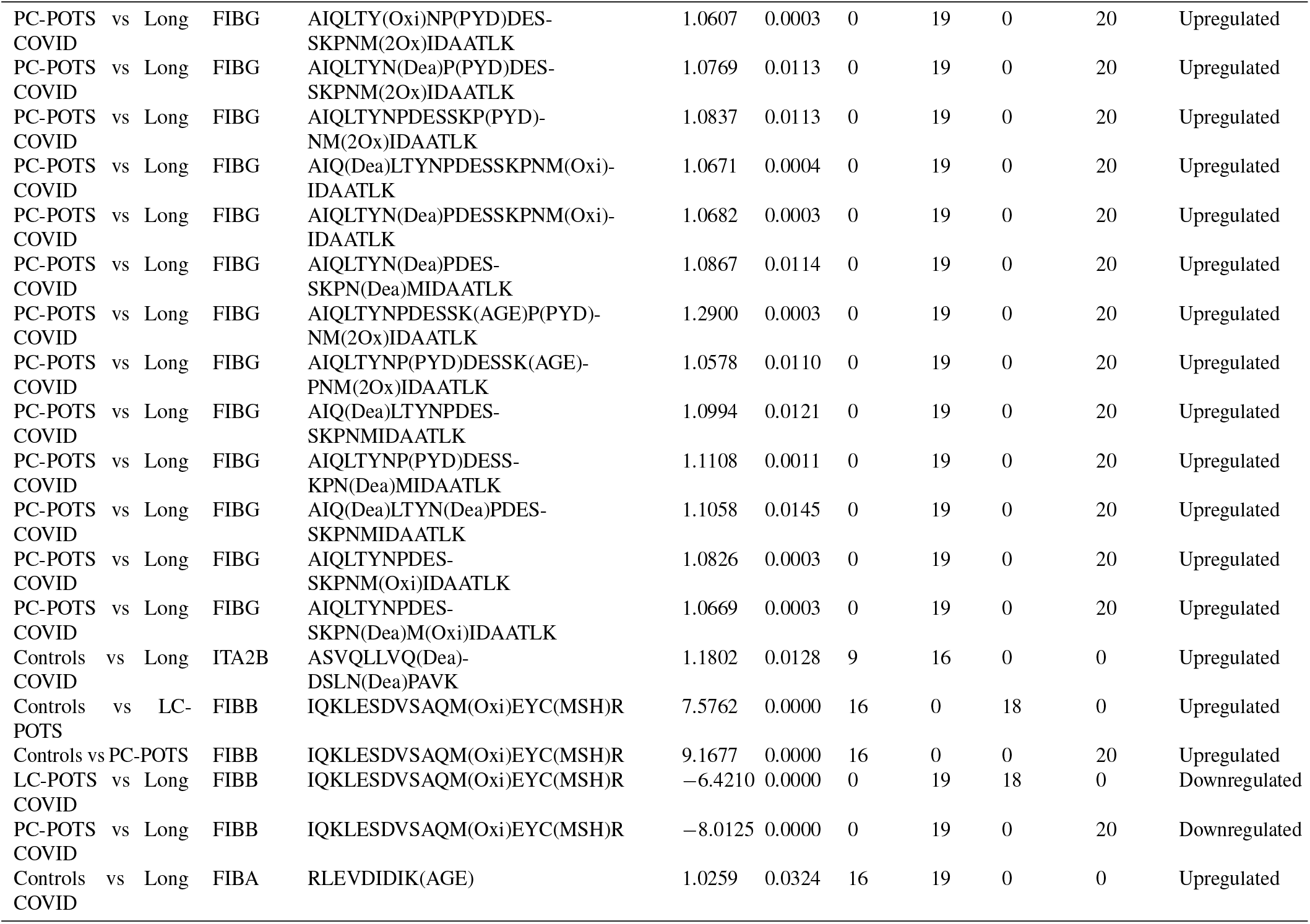
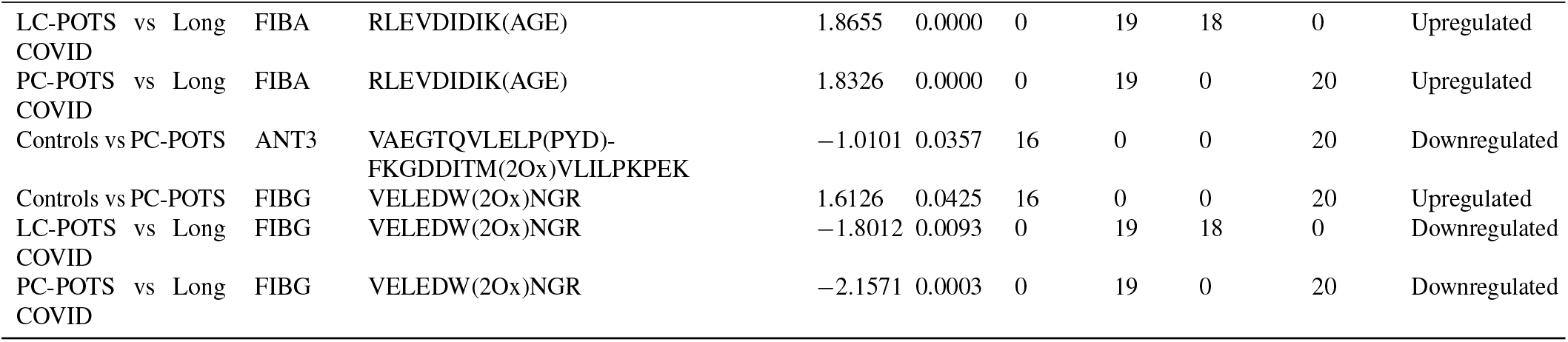
Coagulation Protein Analysis.

**Table S4.2.** Immune Protein Analysis.

| Comparison |  |  | Protein ID | Peptide Modified Sequence | log2FC | Adj. P-Value | Control n | LC n | LC POTS n | POTS n | Direction |
| --- | --- | --- | --- | --- | --- | --- | --- | --- | --- | --- | --- |
| Controls vs LC-POTS |  |  | IGG1 | FN(Dea)WY(Oxi)VDGVEVHNAK | 2.1273 | 0.0043 | 16 | 0 | 18 | 0 | Upregulated |
| Controls vs LC-POTS |  |  | IGG1 | FNW(2Ox)YVDGVEVHNAK | -1.0634 | 0.0032 | 16 | 0 | 18 | 0 | Downregulated |
| Controls vs LC-POTS |  |  | IGG1 | FN(Dea)W(Oxi)YVDGVEVHNAK | 3.0394 | 0.0000 | 16 | 0 | 18 | 0 | Upregulated |
| Controls vs Long COVID |  |  | IGG1 | FN(Dea)WY(Oxi)VDGVEVHN(Dea)AK | -2.4095 | 0.0095 | 14 | 16 | 0 | 0 | Downregulated |
| Controls vs PC-POTS |  |  | IGG1 | FN(Dea)WY(Oxi)VDGVEVHNAK | 2.3740 | 0.0005 | 16 | 0 | 0 | 20 | Upregulated |
| Controls vs PC-POTS |  |  | IGG1 | FNW(Oxi)Y(Oxi)VDGVEVHNAK | 1.3194 | 0.0167 | 16 | 0 | 0 | 20 | Upregulated |
| Controls vs PC-POTS |  |  | IGG1 | FN(Dea)W(Oxi)YVDGVEVHNAK | 3.2874 | 0.0000 | 16 | 0 | 0 | 20 | Upregulated |
| LC-POTS vs Long COVID |  |  | IGG1 | FNW(Oxi)YVDGVEVHNAK | -1.3997 | 0.0193 | 0 | 19 | 18 | 0 | Downregulated |
| LC-POTS vs Long COVID |  |  | IGG1 | FN(Dea)WY(Oxi)VDGVEVHN(Dea)AK | -1.9222 | 0.0093 | 0 | 16 | 18 | 0 | Downregulated |
| LC-POTS vs Long COVID |  |  | IGG1 | FN(Dea)WY(Oxi)VDGVEVHNAK | -2.6590 | 0.0000 | 0 | 19 | 18 | 0 | Downregulated |
| LC-POTS vs Long COVID |  |  | IGG1 | FNW(2Ox)YVDGVEVHNAK | 1.0224 | 0.0071 | 0 | 19 | 18 | 0 | Upregulated |
| LC-POTS vs Long COVID |  |  | IGG1 | FN(Dea)W(Oxi)YVDGVEVHNAK | -3.1352 | 0.0000 | 0 | 19 | 18 | 0 | Downregulated |
| PC-POTS vs Long COVID |  |  | IGG1 | FNW(Oxi)YVDGVEVHNAK | -1.6314 | 0.0022 | 0 | 19 | 0 | 20 | Downregulated |
| PC-POTS vs Long COVID |  |  | IGG1 | FN(Dea)WY(Oxi)VDGVEVHN(Dea)AK | -2.0602 | 0.0015 | 0 | 16 | 0 | 20 | Downregulated |
| PC-POTS vs Long COVID |  |  | IGG1 | FN(Dea)WY(Oxi)VDGVEVHNAK | -2.9057 | 0.0000 | 0 | 19 | 0 | 20 | Downregulated |
| PC-POTS vs Long COVID |  |  | IGG1 | FNW(Oxi)Y(Oxi)VDGVEVHNAK | -1.3773 | 0.0018 | 0 | 19 | 0 | 20 | Downregulated |
| PC-POTS vs Long COVID |  |  | IGG1 | FN(Dea)W(Oxi)YVDGVEVHNAK | -3.3831 | 0.0000 | 0 | 19 | 0 | 20 | Downregulated |
| Controls vs PC-POTS |  |  | IC1 | FQ(Dea)PTLLTLPR | -1.4340 | 0.0167 | 16 | 0 | 0 | 20 | Downregulated |
| Controls vs LC-POTS |  |  | A2MG | IAQW(2Ox)Q(Dea)SFQLEGGLK | 3.1387 | 0.0000 | 16 | 0 | 18 | 0 | Upregulated |
| Controls vs LC-POTS |  |  | A2MG | IAQW(2Ox)QSFQLEGGLK | 2.9169 | 0.0000 | 16 | 0 | 18 | 0 | Upregulated |

Table S4.2 – Continued from previous page
| Comparison | Protein ID | Peptide Modified Sequence | log2FC | Adj. P-Value | Control n | LC n | LC POTS n | POTS n | Direction |
| --- | --- | --- | --- | --- | --- | --- | --- | --- | --- |
| Controls vs LC-POTS | A2MG | IAQ(Dea)W(2Ox)QSFQLEGGLK | 2.9670 | 0.0000 | 16 | 0 | 18 | 0 | Upregulated |
| Controls vs PC-POTS | A2MG | IAQW(Oxi)QSFQLEGGLK | 2.0335 | 0.0167 | 16 | 0 | 0 | 20 | Upregulated |
| Controls vs PC-POTS | A2MG | IAQW(2Ox)Q(Dea)SFQLEGGLK | 3.2468 | 0.0000 | 16 | 0 | 0 | 20 | Upregulated |
| Controls vs PC-POTS | A2MG | IAQW(2Ox)QSFQLEGGLK | 3.0190 | 0.0000 | 16 | 0 | 0 | 20 | Upregulated |
| Controls vs PC-POTS | A2MG | IAQ(Dea)W(2Ox)QSFQLEGGLK | 3.0736 | 0.0000 | 16 | 0 | 0 | 20 | Upregulated |
| LC-POTS vs Long COVID | A2MG | IAQW(Oxi)QSFQLEGGLK | -2.7922 | 0.0001 | 0 | 19 | 18 | 0 | Downregulated |
| LC-POTS vs Long COVID | A2MG | IAQW(2Ox)Q(Dea)SFQLEGGLK | -2.8003 | 0.0000 | 0 | 19 | 18 | 0 | Downregulated |
| LC-POTS vs Long COVID | A2MG | IAQ(Dea)W(2Ox)Q(Dea)SFQLEGGLK | -1.0974 | 0.0259 | 0 | 19 | 18 | 0 | Downregulated |
| LC-POTS vs Long COVID | A2MG | IAQW(2Ox)QSFQLEGGLK | -2.5248 | 0.0001 | 0 | 19 | 18 | 0 | Downregulated |
| LC-POTS vs Long COVID | A2MG | IAQ(Dea)W(2Ox)QSFQLEGGLK | -2.6613 | 0.0000 | 0 | 19 | 18 | 0 | Downregulated |
| PC-POTS vs Long COVID | A2MG | IAQW(Oxi)QSFQLEGGLK | -2.9107 | 0.0000 | 0 | 19 | 0 | 20 | Downregulated |
| PC-POTS vs Long COVID | A2MG | IAQW(2Ox)Q(Dea)SFQLEGGLK | -2.9084 | 0.0000 | 0 | 19 | 0 | 20 | Downregulated |
| PC-POTS vs Long COVID | A2MG | IAQ(Dea)W(2Ox)Q(Dea)SFQLEGGLK | -1.2099 | 0.0054 | 0 | 19 | 0 | 20 | Downregulated |
| PC-POTS vs Long COVID | A2MG | IAQW(2Ox)QSFQLEGGLK | -2.6269 | 0.0000 | 0 | 19 | 0 | 20 | Downregulated |
| PC-POTS vs Long COVID | A2MG | IAQ(Dea)W(2Ox)QSFQLEGGLK | -2.7679 | 0.0000 | 0 | 19 | 0 | 20 | Downregulated |
| Controls vs LC-POTS | CO4A/B | LVN(Dea)GQ(Dea)SHISLSK | 2.5968 | 0.0214 | 10.5 | 0 | 17.5 | 0 | Upregulated |
| Controls vs PC-POTS | CO4A/B | LVN(Dea)GQ(Dea)SHISLSK | 3.0414 | 0.0002 | 10.5 | 0 | 0 | 20 | Upregulated |
| LC-POTS vs Long COVID | CO4A/B | LVN(Dea)GQ(Dea)SHISLSK | -2.6726 | 0.0010 | 0 | 19 | 17.5 | 0 | Downregulated |
| PC-POTS vs Long COVID | CO4A/B | LVN(Dea)GQ(Dea)SHISLSK | -3.1172 | 0.0000 | 0 | 19 | 0 | 20 | Downregulated |
| Controls vs PC-POTS | ITIH4 | N(Dea)M(2Ox)EQ(Dea)FQVSVSVAPNAK | -1.0542 | 0.0149 | 16 | 0 | 0 | 20 | Downregulated |
| Controls vs PC-POTS | ITIH4 | N(Dea)M(Oxi)EQ(Dea)FQVSVSVAPNAK | -1.0384 | 0.0149 | 16 | 0 | 0 | 20 | Downregulated |
| Controls vs PC-POTS | ITIH4 | NM(Oxi)EQ(Dea)FQVSVSVAPNAK | -1.0460 | 0.0160 | 16 | 0 | 0 | 20 | Downregulated |
| Controls vs PC-POTS | ITIH4 | NM(Oxi)EQFQ(Dea)VSVSVAPNAK | -1.0534 | 0.0149 | 16 | 0 | 0 | 20 | Downregulated |

Table S4.2 – Continued from previous page
| Comparison | Protein ID | Peptide Modified Sequence | log2FC | Adj. P-Value | Control n | LC n | LC POTS n | POTS n | Direction |
| --- | --- | --- | --- | --- | --- | --- | --- | --- | --- |
| Controls vs LC-POTS | IGG1 | NQVSLTC(MSH)LVK(AGE) | −3.0791 | 0.0131 | 16 | 0 | 16 | 0 | Downregulated |
| PC-POTS vs LC-POTS | IGG1 | NQVSLTC(MSH)LVK(AGE) | −3.3767 | 0.0142 | 0 | 0 | 16 | 20 | Downregulated |
| Controls vs PC-POTS | IGG1 | NTLY(Oxi)LNMNSLRP(PYD)-EDTAVYYC(MSH)AR | 1.2630 | 0.0092 | 16 | 0 | 0 | 20 | Upregulated |
| Controls vs Long COVID | IGG1 | P(PYD)KDTLM(2Ox)ISRTPEVTCVVVDVSHEDPEVK | 1.2217 | 0.0450 | 16 | 19 | 0 | 0 | Upregulated |
| LC-POTS vs Long COVID | IGG1 | P(PYD)KDTLM(2Ox)ISRTPEVTCV-VVDVSHEDPEVK | 2.2216 | 0.0000 | 0 | 19 | 18 | 0 | Upregulated |
| PC-POTS vs Long COVID | IGG1 | P(PYD)KDTLM(2Ox)ISRTPEVTCV-VVDVSHEDPEVK | 1.7612 | 0.0003 | 0 | 19 | 0 | 20 | Upregulated |
| Controls vs PC-POTS | A2MG | QFSFPLSSEPFQ(Dea)GSYK | 1.0692 | 0.0249 | 16 | 0 | 0 | 20 | Upregulated |
| Controls vs PC-POTS | CO3 | SEETKEN(Dea)EGFTVTAEGK | 1.0172 | 0.0261 | 15 | 0 | 0 | 7 | Upregulated |
| Controls vs LC-POTS | IGG1 | THTC(MSH)PPCPAPELLG-GPSVFLFPPKPK | 2.7867 | 0.0001 | 14 | 0 | 18 | 0 | Upregulated |
| Controls vs PC-POTS | IGG1 | THTC(MSH)PPCPAPELLG-GPSVFLFPPKPK | 2.6974 | 0.0001 | 14 | 0 | 0 | 20 | Upregulated |
| LC-POTS vs Long COVID | IGG1 | THTC(MSH)PPCPAPELLG -GPSVFLFPPKPK | −1.8519 | 0.0031 | 0 | 19 | 18 | 0 | Downregulated |
| PC-POTS vs Long COVID | IGG1 | THTC(MSH)PPCPAPELLG-GPSVFLFPPKPK | −1.7625 | 0.0011 | 0 | 19 | 0 | 20 | Downregulated |
| Controls vs PC-POTS | IGG1 | VVSVLTVLHQDW(Oxi)LN(Dea)GK | 1.1606 | 0.0149 | 13 | 0 | 0 | 20 | Upregulated |
| LC-POTS vs Long COVID | IGG1 | VVSVLTVLHQDW(Oxi)LN(Dea)GK | −1.0605 | 0.0352 | 0 | 19 | 18 | 0 | Downregulated |
| PC-POTS vs Long COVID | IGG1 | VVSVLTVLHQDW(Oxi)LN(Dea)GK | −1.3110 | 0.0035 | 0 | 19 | 0 | 20 | Downregulated |
| Controls vs PC-POTS | CO3 | W(3Ox)LNEQRYYYGGYGSTQ(Dea)-ATFMVFQALAQYQK | −1.0699 | 0.0149 | 16 | 0 | 0 | 20 | Downregulated |
| Controls vs PC-POTS | CO3 | YYGGGYGSTQ(Dea)ATFMVFQALAQYQK | −1.0326 | 0.0169 | 15 | 0 | 0 | 19 | Downregulated |

**Table S4.3.** Metabolic Protein Analysis.

| Comparison | Protein ID | Peptide Modified Sequence | log2FC | Adj. P-Value | Control n | LC n | LC POTS n | POTS n | Direction |
| --- | --- | --- | --- | --- | --- | --- | --- | --- | --- |
| Controls vs PC-POTS | ALBU | DDN(Dea)PN(Dea)LPR(AGE)-LVRPEVDVMCTAFHDNEETFLK | -1.0002 | 0.0003 | 16 | 0 | 0 | 20 | Downregulated |
| Controls vs PC-POTS | ALBU | DLGEENFK | -1.3517 | 0.0302 | 16 | 0 | 0 | 20 | Downregulated |
| PC-POTS vs Long COVID | ALBU | DLGEENFK | 1.3559 | 0.0087 | 0 | 19 | 0 | 20 | Upregulated |
| Controls vs PC-POTS | ALBU | EFNAETFTFHADI-C(MSH)TLSEKER(AGE) | -1.4501 | 0.0426 | 16 | 0 | 0 | 20 | Downregulated |
| PC-POTS vs Long COVID | ALBU | EFN(Dea)AETFTFHADI-C(MSH)TLSEKER(AGE) | 1.0219 | 0.0498 | 0 | 19 | 0 | 20 | Upregulated |
| Controls vs LC-POTS | APOA1 | LLDN(Dea)W(2Ox)DSVTSTFSK | 3.6468 | 0.0000 | 16 | 0 | 18 | 0 | Upregulated |
| Controls vs LC-POTS | APOA1 | LLDNW(Oxi)DSVTSTFSK | 3.3183 | 0.0000 | 16 | 0 | 18 | 0 | Upregulated |
| Controls vs LC-POTS | APOA1 | LLDNW(2Ox)DSVTSTFSK | 3.8719 | 0.0000 | 16 | 0 | 18 | 0 | Upregulated |
| Controls vs LC-POTS | APOA1 | LLDN(Dea)W(Oxi)DSVTSTFSK | 2.7881 | 0.0000 | 16 | 0 | 18 | 0 | Upregulated |
| Controls vs PC-POTS | APOA1 | LLDN(Dea)W(2Ox)DSVTSTFSK | 3.6733 | 0.0000 | 16 | 0 | 0 | 20 | Upregulated |
| Controls vs PC-POTS | APOA1 | LLDNW(Oxi)DSVTSTFSK | 3.3715 | 0.0000 | 16 | 0 | 0 | 20 | Upregulated |
| Controls vs PC-POTS | APOA1 | LLDNW(2Ox)DSVTSTFSK | 3.9021 | 0.0000 | 16 | 0 | 0 | 20 | Upregulated |
| Controls vs PC-POTS | APOA1 | LLDN(Dea)W(Oxi)DSVTSTFSK | 2.7370 | 0.0000 | 16 | 0 | 0 | 20 | Upregulated |
| LC-POTS vs Long COVID | APOA1 | LLDN(Dea)W(2Ox)DSVTSTFSK | -3.8376 | 0.0000 | 0 | 19 | 18 | 0 | Downregulated |
| LC-POTS vs Long COVID | APOA1 | LLDNW(Oxi)DSVTSTFSK | -3.2224 | 0.0000 | 0 | 19 | 18 | 0 | Downregulated |
| LC-POTS vs Long COVID | APOA1 | LLDNW(2Ox)DSVTSTFSK | -3.8403 | 0.0000 | 0 | 19 | 18 | 0 | Downregulated |
| LC-POTS vs Long COVID | APOA1 | LLDN(Dea)W(Oxi)DSVTSTFSK | -3.4010 | 0.0000 | 0 | 19 | 18 | 0 | Downregulated |
| PC-POTS vs Long COVID | APOA1 | LLDN(Dea)W(2Ox)DSVTSTFSK | -3.8641 | 0.0000 | 0 | 19 | 0 | 20 | Downregulated |
| PC-POTS vs Long COVID | APOA1 | LLDNW(Oxi)DSVTSTFSK | -3.2756 | 0.0000 | 0 | 19 | 0 | 20 | Downregulated |
| PC-POTS vs Long COVID | APOA1 | LLDNW(2Ox)DSVTSTFSK | -3.8704 | 0.0000 | 0 | 19 | 0 | 20 | Downregulated |

Table S4.3 – Continued from previous page
| Comparison | Protein ID | Peptide Modified Sequence | log2FC | Adj. P-Value | Control n | LC n | LC POTS n | POTS n | Direction |
| --- | --- | --- | --- | --- | --- | --- | --- | --- | --- |
| PC-POTS vs Long COVID | APOA1 | LLDN(Dea)W(Oxi)DSVTSTFSK | −3.3499 | 0.0000 | 0 | 19 | 0 | 20 | Downregulated |
| Controls vs LC-POTS | HBB | LLVVYPWTQ(Dea)R | −1.0031 | 0.0374 | 16 | 0 | 18 | 0 | Downregulated |
| Controls vs Long COVID | HBB | LLVVYPWTQ(Dea)R | −1.2375 | 0.0080 | 16 | 19 | 0 | 0 | Downregulated |
| Controls vs Long COVID | HBA | M(Oxi)FLSFPTTK | −1.1655 | 0.0431 | 16 | 19 | 0 | 0 | Downregulated |
| Controls vs PC-POTS | ALBU | P(PYD)LLEKSHCIAEVENDEM-PADLPSLAADFVESK | −1.0333 | 0.0373 | 16 | 0 | 0 | 20 | Downregulated |
| Controls vs Long COVID | ALBU | RP(PYD)CFSALEVDETYVVPK | −1.2540 | 0.0117 | 16 | 19 | 0 | 0 | Downregulated |
| LC-POTS vs Long COVID | ALBU | RP(PYD)CFSALEVDETYVVPK | −1.1361 | 0.0286 | 0 | 19 | 18 | 0 | Downregulated |
| PC-POTS vs Long COVID | ALBU | RP(PYD)CFSALEVDETYVVPK | −1.1836 | 0.0137 | 0 | 19 | 0 | 20 | Downregulated |
| Controls vs PC-POTS | APOB | Y(Oxi)ENYELTLK | 1.4558 | 0.0425 | 16 | 0 | 0 | 20 | Upregulated |

## References

[1] D Dharmindra Dulal, A Maraey, H Elsharnoby, P Chacko, and B Grubb. Impact of COVID-19 pandemic on the incidence and prevalence of postural orthostatic tachycardia syndrome. Eur Heart J Qual Care Clin Outcomes, 11(5): 698–704, 2025. doi: 10.1093/ehjqcco/qcae111.

[2] S J Yong. Long COVID or post-COVID-19 syndrome: putative pathophysiology, risk factors, and treatments. Infect Dis, 53(10):737–754, 2021. doi: 10.1080/23744235.2021.1924397.

[3] Hannah Mandel, Yun J Yoo, Andrea J Allen, Sajjad Abedian, Zoe Verzani, Elizabeth W Karlson, Lawrence C Kleinman, Praveen C Mudumbi, Carlos R Oliveira, Jennifer A Muszynski, Rachel S Gross, Thomas W Carton, C Kim, Emily Taylor, Heekyong Park, Jasmin Divers, J Daniel Kelly, Jonathan Arnold, Carol Reynolds Geary, Chengxi Zang, Kelan G Tantisira, Kyung E Rhee, Michael Koropsak, Sindhu Mohandas, Andrew Vasey, Abu Saleh Mohammad Mosa, Melissa Haendel, Christopher G Chute, Shawn N Murphy, Lisa O’Brien, Jacqueline Szmuszkovicz, Nicholas Guthe, Jorge L Santana, Aliva De, Amanda L Bogie, Katia C Halabi, Lathika Mohanraj, Patricia A Kinser, Samuel E Packard, Katherine R Tuttle, Kathryn Hirabayashi, Rainu Kaushal, Emily Pfaff, Mark G Weiner, Lorna E Thorpe, and Richard A Moffitt. Long COVID Incidence Proportion in Adults and Children Between 2020 and 2024: An Electronic Health Record-Based Study From the RECOVER Initiative. Clinical infectious diseases : an official publication of the Infectious Diseases Society of America, 80(6):1247–1261, 7 2025. ISSN 1537-6591. doi: 10.1093/cid/ciaf046.

[4] Bozkurt B, Das SR, Addison D, Gupta A, Jneid H, Khan SS, Koromia GA, Kulkarni PA, LaPoint K, Lewis EF, Michos ED, Peterson PN, Turagam MK, Wang TY, and Yancy CW. 2022 AHA/ACC key data elements and definitions for cardiovascular and noncardiovascular complications of COVID-19. Journal of the American College of Cardiology, 80(4):388–465, 2022. doi: 10.1016/j.jacc.2022.03.355.

[5] A. Fedorowski. Postural orthostatic tachycardia syndrome: clinical presentation, aetiology and management. Journal of Internal Medicine, 285(4):352–366, 4 2019. ISSN 0954-6820. doi: 10.1111/joim.12852.

[6] S Vernino, KM Bourne, LE Stiles, BP Grubb, A Fedorowski, JM Stewart, AC Arnold, LA Pace, J Axelsson, JR Boris, JP Moak, BP Goodman, KR Chémali, TH Chung, DS Goldstein, A Diedrich, MG Miglis, MM Cortez, AJ Miller, R Freeman, I Biaggioni, PC Rowe, RS Sheldon, CA Shibao, DM Systrom, GA Cook, TA Doherty, HI Abdallah, A Darbari, and SR Raj. Postural orthostatic tachycardia syndrome (POTS): State of the science and clinical care from a 2019 National Institutes of Health Expert Consensus Meeting - Part 1. Auton Neurosci, 235:102828, 2021. doi: 10.1016/j.autneu.2021.102828.

[7] SR Raj, KM Bourne, LE Stiles, MG Miglis, MM Cortez, AJ Miller, R Freeman, I Biaggioni, PC Rowe, RS Sheldon, CA Shibao, A Diedrich, DM Systrom, GA Cook, TA Doherty, HI Abdallah, BP Grubb, A Fedorowski, JM Stewart, AC Arnold, LA Pace, J Axelsson, JR Boris, JP Moak, BP Goodman, K R Chémali, TH Chung, DS Goldstein, A Darbari, and S Vernino. Postural orthostatic tachycardia syndrome (POTS): Priorities for POTS care and research from a 2019 National Institutes of Health Expert Consensus Meeting - Part 2. Auton Neurosci, 235:102836, 2021. doi: 10.1016/j.autneu.2021.102836.

[8] B H Shaw, L E Stiles, K Bourne, EA Green, CA Shibao, LE Okamoto, EM Garland, A Gamboa, A Diedrich, V Raj, RS Sheldon, I Biaggioni, D Robertson, and SR Raj. The face of postural tachycardia syndrome - insights from a large cross-sectional online community-based survey. J Intern Med, 286(4):438–448, 2019. doi: 10.1111/joim.12895.

[9] N L DePace and J Colombo. Long-COVID syndrome and the cardiovascular system: A review of neurocardiologic effects on multiple systems. Curr Cardiol Rep, 24:1711–1726, 2022. doi: 10.1007/s11886-022-01786-2.

[10] Buonsenso D, Piazza M, Boner AL, and Bellanti JA. Long COVID: A proposed hypothesis-driven model of viral persistence for the pathophysiology of the syndrome. Allergy Asthma Proc., 43(3):187–193, 2022. doi: 10.2500/aap.2022.43.220018.

[11] Kavi Lesley. Br J Gen Pract. Br J Gen Pract, 72(714): 8–9, 2022. doi: 10.3399/bjgp22X718037.

[12] M Johansson, H Yan, C Welinder, and others. Plasma proteomic profiling in postural orthostatic tachycardia syndrome (POTS) reveals new disease pathways. Sci Rep, 12:20051, 2022. doi: 10.1038/s41598-022-24729-x.

[13] E Pretorius, C Venter, G J Laubscher, P J Lourens, J Steenkamp, and DB Kell. Prevalence of readily detected amyloid blood clots in ‘unclotted’ Type 2 Diabetes Mellitus and COVID-19 plasma: a preliminary report. Cardiovasc Diabetol, 19:193, 2020. doi: 10.1186/s12933-020-01165-7.

[14] E Pretorius, M Vlok, C Venter, JA Bezuidenhout, GJ Laubscher, J Steenkamp, and DB Kell. Persistent clotting protein pathology in Long COVID/Post-Acute Sequelae of COVID-19 (PASC) is accompanied by increased levels of antiplasmin. Cardiovasc Diabetol, 20:172, 2021. doi: 10.1186/s12933-021-01359-7.

[15] Douglas B. Kell, Gert Jacobus Laubscher, and Etheresia Pretorius. A central role for amyloid fibrin microclots in long COVID/PASC: origins and therapeutic implications. Biochemical Journal, 479(4):537–559, 2 2022. ISSN 0264-6021. doi: 10.1042/BCJ20220016.

[16] Alain R. Thierry, Tom Usher, Cynthia Sanchez, Simone Turner, Chantelle Venter, Brice Pastor, Maxine Waters, Anel Thompson, Alexia Mirandola, Ekaterina Pisareva, Corinne Prevostel, Gert J. Laubscher, Douglas B. Kell, and Etheresia Pretorius. Circulating Microclots Are Structurally Associated With Neutrophil Extracellular Traps and Their Amounts Are Elevated in Long COVID Patients. Journal of Medical Virology, 97(10), 10 2025. ISSN 0146-6615. doi: 10.1002/jmv.70613.

[17] Bisaccia G, Ricci F, Recce V, Serio A, Iannetti G, Chahal AA, Ståhlberg M, Khanji MY, Fedorowski A, and Gallina S. Post-Acute Sequelae of COVID-19 and Cardiovascular Autonomic Dysfunction: What Do We Know? Journal of Cardiovascular Development and Disease, 8(11):156, 2021. doi: 10.3390/jcdd8110156.

[18] M Johansson, M Ståhlberg, M Runold, M Nygren-Bonnier, J Nilsson, B Olshansky, J Bruchfeld, and A Fedorowski. Long-Haul Post-COVID-19 Symptoms Presenting as a Variant of Postural Orthostatic Tachycardia Syndrome: The Swedish Experience. JACC Case Rep, 3 (4):573–580, 2021. doi: 10.1016/j.jaccas.2021.01.009.

[19] K Kanjwal, S Jamal, A Kichloo, and BP Grubb. New-onset Postural Orthostatic Tachycardia Syndrome Following Coronavirus Disease 2019 Infection. J Innov Card Rhythm Manag, 11(11):4302–4304, 2020. doi: 10.19102/icrm.2020.111102.

[20] W T 3rd Gunning, S Khan, JW Spatafore, BL Karabin, and BP Grubb. Postural orthostatic tachycardia syndrome in post-COVID-19 long-hauler patients is associated with platelet storage pool deficiency. Front Med (Lausanne), 12:1560120, 2025. doi: 10.3389/fmed.2025.1560120.

[21] Douglas B. Kell, Muhammed Asad Khan, Binita Kane, Gregory Y. H. Lip, and Etheresia Pretorius. Possible Role of Fibrinaloid Microclots in Postural Orthostatic Tachycardia Syndrome (POTS): Focus on Long COVID. Journal of Personalized Medicine, 14(2):170, 1 2024. ISSN 2075-4426. doi: 10.3390/jpm14020170.

[22] A Kruger, M Vlok, S Turner, and others. Proteomics of fibrin amyloid microclots in long COVID/post-acute sequelae of COVID-19 (PASC) shows many entrapped pro-inflammatory molecules that may also contribute to a failed fibrinolytic system. Cardiovasc Diabetol, 21:190, 2022. doi: 10.1186/s12933-022-01623-4.

[23] Callum Thomas, Mark A. Faghy, Corinna Chidley, Bethan E. Phillips, Thomas Bewick, and Ruth E Ashton. Blood Biomarkers of Long COVID: A Systematic Review. Molecular Diagnosis & Therapy, 28(5):537–574, 9 2024. ISSN 1177-1062. doi: 10.1007/s40291-024-00731-z.

[24] H Dutta and N Jain. Post-translational modifications and their implications in cancer. Front Oncol, 13:1240115, 2023. doi: 10.3389/fonc.2023.1240115.

[25] F Nencini, A Bettiol, FR Argento, S Borghi, E Giurranna, G Emmi, D Prisco, N Taddei, C Fiorillo, and M Becatti. Post-translational modifications of fibrinogen: implications for clotting, fibrin structure and degradation. Mol Biomed, 5(1):45, 2024. doi: 10.1186/s43556-024-00214-x.

[26] Marie Mikuteit, Svetlana Baskal, Sandra Klawitter, Alexandra Dopfer-Jablonka, Georg M N Behrens, Frank Müller, Dominik Schröder, Frank Klawonn, Sandra Steffens, and Dimitrios Tsikas. Amino acids, post-translational modifications, nitric oxide, and oxidative stress in serum and urine of long COVID and ex COVID human subjects. Amino acids, 55(9):1173–1188, 9 2023. ISSN 1438-2199. doi: 10.1007/s00726-023-03305-1.

[27] DB Kell and E Pretorius. Proteomic evidence for amyloidogenic cross-seeding in fibrinaloid microclots. Int J Mol Sci, 25(19):10809, 2024. doi: 10.3390/ijms251910809.

[28] Douglas B. Kell and Etheresia Pretorius. The Proteome Content of Blood Clots Observed Under Different Conditions: Successful Role in Predicting Clot Amyloid(ogenicity). Molecules, 30(3):668, 2 2025. ISSN 1420-3049. doi: 10.3390/molecules30030668.

[29] Douglas B. Kell, Karen M. Doyle, J. Enrique Salcedo-Sora, Alakendu Sekhar, Melanie Walker, and Etheresia Pretorius. AmyloGram reveals amyloidogenic potential in stroke thrombus proteomes. Biochemical Journal, 482 (22):1689–1706, 11 2025. ISSN 0264-6021. doi: 10.1042/BCJ20253317.

[30] Justine M. Grixti, Arun Chandran, Jan-Hendrik Pretorius, Melanie Walker, Alakendu Sekhar, Etheresia Pretorius, and Douglas B. Kell. Amyloid Presence in Acute Ischemic Stroke Thrombi: Observational Evidence for Fibrinolytic Resistance. Stroke, 56(7), 7 2025. ISSN 0039-2499. doi: 10.1161/STROKEAHA.124.050033.

[31] N Sheikh, S Ranada, K Kogut, and others. Exploring the Refractory Period of an Active Stand in Females With Initial Orthostatic Hypotension. JACC, 77(25):3228–3229, 2021. doi: 10.1016/j.jacc.2021.04.068.

[32] Nasia A Sheikh, Shaun Ranada, Matthew Lloyd, Dallan McCarthy, Karolina Kogut, Kate M Bourne, Juliana G Jorge, Lucy Y Lei, Robert S Sheldon, Derek V Exner, Aaron A Phillips, Mary Runté, and Satish R Raj. Lower body muscle preactivation and tensing mitigate symptoms of initial orthostatic hypotension in young females. Heart rhythm, 19(4):604–610, 4 2022. ISSN 1556-3871. doi: 10.1016/j.hrthm.2021.12.030.

[33] SR Raj, JC Guzman, P Harvey, L Richer, R Schondorf, C Seifer, N Thibodeau-Jarry, and RS Sheldon. Canadian Cardiovascular Society Position Statement on Postural Orthostatic Tachycardia Syndrome (POTS) and Related Disorders of Chronic Orthostatic Intolerance. Can J Cardiol, 36(3):357–372, 2020. doi: 10.1016/j.cjca.2019.12.024.

[34] Simone Turner, Gert Jacobus Laubscher, M Asad Khan, Douglas B. Kell, and Etheresia Pretorius. Accelerating discovery: A novel flow cytometric method for detecting fibrin(ogen) amyloid microclots using long COVID as a model. Heliyon, 9(9):e19605, 9 2023. ISSN 24058440. doi: 10.1016/j.heliyon.2023.e19605.

[35] F Nencini, E Giurranna, S Borghi, N Taddei, C Fiorillo, and M Becatti. Fibrinogen oxidation and thrombosis: shaping structure and function. Antioxidants (Basel), 14 (4):390, 2025. doi: 10.3390/antiox14040390.

[36] A Twarda-Clapa, A Olczak, A M Białkowska, and M Koziołkiewicz. Advanced glycation end-products (AGEs): Formation, chemistry, classification, receptors, and diseases related to AGEs. Cells, 11(8):1312, 2022. doi: 10.3390/cells11081312.

[37] Bychkova AV, Vasilyeva AD, Bugrova AE, Indeykina MI, Kononikhin AS, Nikolaev EN, Konstantinova ML, and Rosenfeld MA. Oxidation-induced modification of the fibrinogen polypeptide chains. Dokl Biochem Biophys, 474 (1):173–177, 2017. doi: 10.1134/S1607672917030115.

[38] E Pretorius, C Venter, JM Nunes, AR Thierry, and DB Kell. Inflammatory Triggers, Cell Death, Membrane Damage and Lipid Asymmetry That Shape Procoagulant Surfaces for Amyloidogenic Microclotting. 10.20944/preprints202510.1460.v1, 2025.

[39] T Stief, A Aab, and N Heimburger. Oxidative inactivation of purified human alpha-2-antiplasmin, antithrombin III, and C1-inhibitor. Thromb Res, 49(6):581–589, 1988. doi: 10.1016/0049-3848(88)90255-1.

[40] B Yan, D Hu, S Knowles, and J Smith. Probing chemical and conformational differences in the resting and active conformers of platelet integrin alpha(IIb)beta(3). J Biol Chem, 275(10):7249–7260, 2000.

[41] Erin E West, Martin Kolev, and Claudia Kemper. Complement and the Regulation of T Cell Responses. Annual review of immunology, 36:309–338, 4 2018. ISSN 1545-3278. doi: 10.1146/annurev-immunol-042617-053245.

[42] Daniel Elieh Ali Komi, Farzaneh Shafaghat, Petri T. Kovanen, and Seppo Meri. Mast cells and complement system: Ancient interactions between components of innate immunity. Allergy, 75(11):2818–2828, 11 2020. ISSN 0105-4538. doi: 10.1111/all.14413.

[43] Ritsuko Kohno, David S. Cannom, Brian Olshansky, Shijun Cindy Xi, Darshan Krishnappa, Wayne O. Adkisson, Faye L. Norby, Artur Fedorowski, and David G. Benditt. Mast Cell Activation Disorder and Postural Orthostatic Tachycardia Syndrome: A Clinical Association. Journal of the American Heart Association, 10(17), 9 2021. ISSN 2047-9980. doi: 10.1161/JAHA.121.021002.

[44] J Vandooren and Y Itoh. Alpha-2-macroglobulin in inflammation, immunity and infections. Front Immunol, 12: 803244, 2021. doi: 10.3389/fimmu.2021.803244.

[45] J Verhelst and Y Impens. Alpha-2-macroglobulin in inflammation, immunity and infections. Front Immunol, 12: 803244, 2021. doi: 10.3389/fimmu.2021.803244.

[46] D Jing, J Jin, Z Mei, Q Zhu, Y Lu, and X Wang. Effects of Helicobacter pylori infection and interleukin 6 on the expression of ITIH4 in human gastric cancer cells. Transl Cancer Res, 9:4656–4665, 2020. doi: 10.21037/tcr-20-1766.

[47] Chloé Chevalier, Arsênio Rodrigues Oliveira, Valentin Blanchard, Cédric Le May, Bertrand Cariou, Samy Hadjadj, and Mikaël Croyal. Post-translational modifications of apolipoproteins as promising biomarkers for diabetes-related cardiovascular diseases: A comprehensive review. Diabetes & Metabolism, 51(5):101683, 9 2025. ISSN 12623636. doi: 10.1016/j.diabet.2025.101683.

[48] Borén J, Packard CJ, and Binder CJ. Apolipoprotein B-containing lipoproteins in atherogenesis. Nat Rev Cardiol, 22:399–413, 2025. doi: 10.1038/s41569-024-01111-0.

[49] M Yang, R Liu, S Li, Y Luo, Y Zhang, L Zhang, D Liu, Y Wang, Z Xiong, G Boden, S Chen, L Li, and G Yang. Zinc-α2-glycoprotein is associated with insulin resistance in humans and is regulated by hyperglycemia, hyperinsulinemia, or liraglutide administration: cross-sectional and interventional studies in normal subjects, insulin-resistant subjects, and subjects with newly diagnosed diabetes. Diabetes Care, 36(5):1074–1082, 2013. doi: 10.2337/dc12-0940.

[50] È Navarro-Masip, DM Selva, C Hernández, A Ciudin, B Salinas-Roca, J Cabrera-Serra, R Simó, and A Lecube. Acute downregulation of zinc α2-glycoprotein: Evidence from human and HepG2 cell studies. Int J Mol Sci, 26 (12):5438, 2025. doi: 10.3390/ijms26125438.

[51] Carolyn T Bramante, John B Buse, David M Liebovitz, Jacinda M Nicklas, Michael A Puskarich, Ken Cohen, Hrishikesh K Belani, Blake J Anderson, Jared D Huling, Christopher J Tignanelli, Jennifer L Thompson, Matthew Pullen, Esteban Lemus Wirtz, Lianne K Siegel, Jennifer L Proper, David J Odde, Nichole R Klatt, Nancy E Sherwood, Sarah M Lindberg, Amy B Karger, Kenneth B Beckman, Spencer M Erickson, Sarah L Fenno, Katrina M Hartman, Michael R Rose, Tanvi Mehta, Barkha Patel, Gwendolyn Griffiths, Neeta S Bhat, Thomas A Murray, David R Boulware, and COVID-OUT Study Team. Outpatient treatment of COVID-19 and incidence of post-COVID-19 condition over 10 months (COVID-OUT): a multicentre, randomised, quadruple-blind, parallel-group, phase 3 trial. The Lancet. Infectious diseases, 23(10):1119–1129, 10 2023. ISSN 1474-4457. doi: 10.1016/S1473-3099(23)00299-2.

